# Untangling Overlapping Barcodes in Image-Based Spatial Genomics

**DOI:** 10.1101/2025.06.10.658913

**Authors:** Jonathan A. White, Chuqi Lu, Lincoln Ombelets, Long Cai

**Affiliations:** Division of Biology and Biological Engineering, California Institute of Technology, Pasadena, CA, USA; Division of Chemistry and Chemical Engineering, California Institute of Technology, Pasadena, CA, USA

**Keywords:** Spatial Genomics, Super-Resolution Microscopy

## Abstract

Difficulty in resolving spatially overlapping barcodes is a major bottleneck for imaging-based spatial genomics methods. Here, we present an approach for untangling spatially overlapping barcodes by using strong encoding and global optimization to reduce spurious solutions resulting from recombinations of barcode signals. We demonstrate experimentally that cellular regions with average local densities of 127 barcodes per ***µm***^***2***^ can be decoded with an estimated FDR of less than **4%**, enabling a new type of super-resolution microscopy by coding.

## 1 Main Text

Spatial genomics technologies measure gene expression of cells in their native tissue context. Imaging-based spatial transcriptomics, such as seqFISH [1], MERFISH [2], hybISS [3], and STARMAP [4] build on single-molecule fluorescent in situ hybridization (smFISH) [5, 6]. In FISH-based methods, sequential cycles of probe hybridization, imaging, and probe stripping (hybridization cycles) generate many images containing barcodes that uniquely identify RNA species. In seqFISH, the images are partitioned into blocks that each represent a symbol of an error-correcting code. We say that each image in a symbol block is a different pseudo-color of the symbol block image: all dots found in a block represent a symbol with a value determined by their pseudo-color number. Each gene is encoded by a unique sequence of symbols called a codeword. Barcodes, the physical manifestation of codewords, appear in seqFISH data as an aligned sequence of signals (dots) in different symbol blocks (Fig. 1A-B)[7–9].

**Fig. 1.**
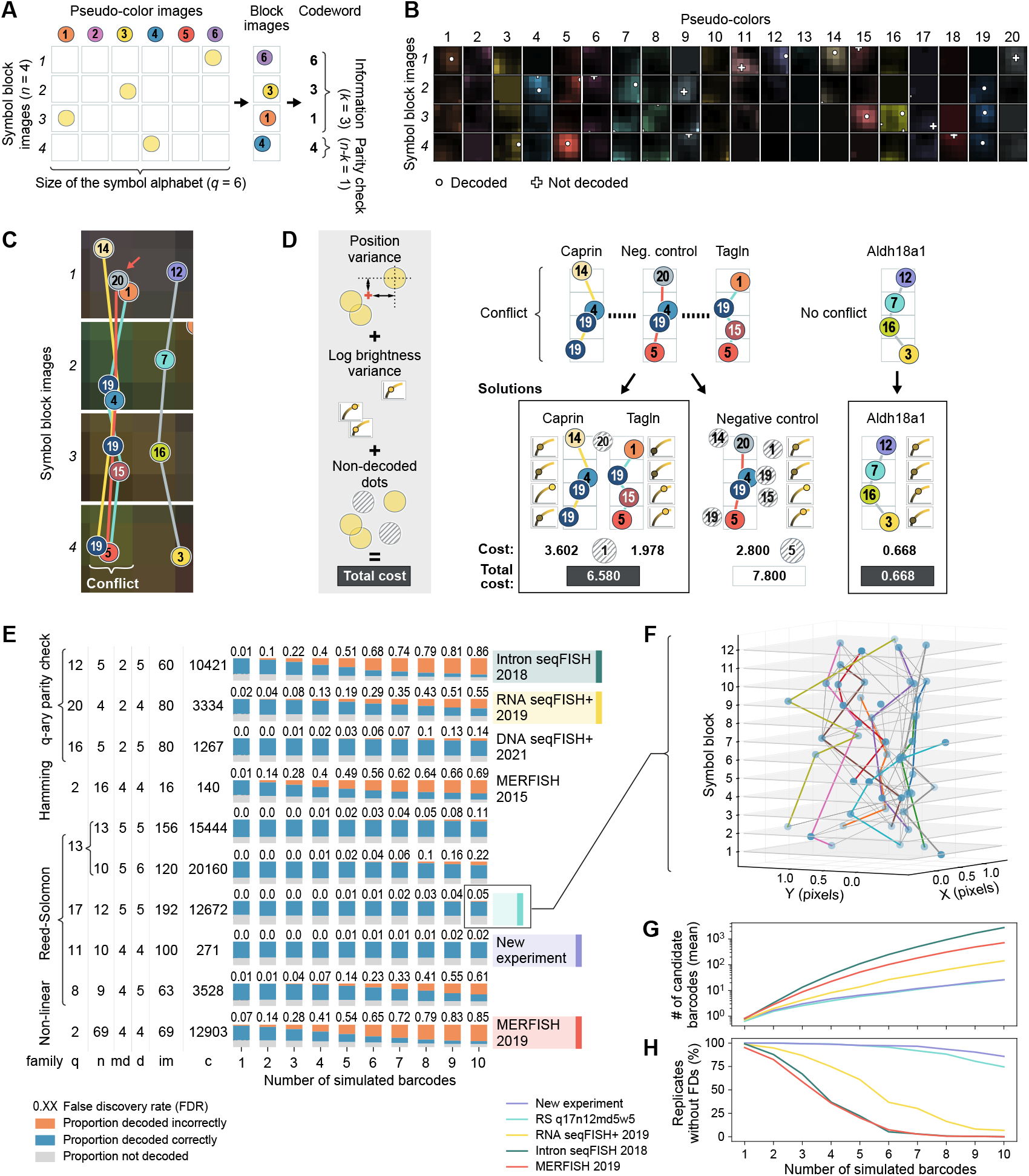
Untangling spatially overlapping barcodes A) An illustration of how seqFISH encodes molecular identity. SeqFISH images are partitioned into symbol blocks (rows) in which symbol values (also called pseudo-colors, shown as columns) transmitted by a barcoded molecule are one-hot encoded. Individual molecules transmit signals that appear as a diffraction-limited dot of one pseudo-color in a symbol block. Sequences of spatially aligned dots of varying pseudo-colors in different symbol blocks represent a sequence of symbols called a codeword that encodes the identity of the barcoded molecule. Codewords are composed of information symbols, which alone specify molecular identity, and parity check symbols, which give the encoding robustness to error. Panels B-D show a decoding example of three superimposed barcodes from the 2019 seqFISH+ experiment. B) An aligned region of interest showing all 80 images in the experiment containing three superimposed barcodes and other fluorescent signals. Images are organized by symbol block (rows) and pseudo-color (columns). White circles are overlaid at the coordinates of fit dots that were decoded. White crosses are overlaid at the coordinates of fit dots that were not decoded. Panel C) shows candidate barcodes identified from the dots in panel B. Images of each symbol block show pixels pseudo-colored according to the aligned fluorescent signal of each pseudo-colored image. The locations of dots centered in the region of interest are marked by circles of their pseudo-color and labeled with the number of their pseudo-color. Candidate barcodes are connected by lines of a color unique to each barcode. The Aldh18a1 barcode (gray) is unambiguous, but the Tagln (blue), Caprin (yellow), and negative control (red) barcodes are conflicting. In this case, it appears that a non-specific dot, pseudo-color 20 in symbol block 1, introduces a spurious negative control barcode candidate straddling the Tagln and Caprin barcodes. D) To resolve the conflicts and find the most likely solution, we transform the network of conflicting barcodes into a graph where each node represents a barcode and edges represent conflicts. Feasible solutions cannot contain barcodes connected by an edge. Each solution is scored by summing the costs of the barcodes it contains and the costs of not decoding each dot that is not decoded. The lowest cost solution is chosen using integer programming. E) Decoding simulated data of numerous barcodes in a 1-pixel (100×100nm) area with our decoding algorithm determines the relative robustness of different error-correcting codes to spurious solutions arising from superimposed barcodes. The dendrogram on the left groups codes by their family first, then various parameters: q, symbol alphabet size; n, number of symbols; md, minimum Hamming distance between any pair of codewords; d, the number of dots are in each barcode; im, the number of images required for an experimental implementation; and c, the number of codewords used from the code. Each condition used 500 replicates. F) A simulation replicate containing 10 superimposed barcodes encoded by the q17 n12 Reed-Solomon code used to produce panel D was correctly decoded. The correct decoded barcodes are shown in colored paths. The gray paths are the candidate barcodes that are not decoded. In this simulation replicate, the algorithm correctly determined all the simulated genes. G-H) Panels G and H, respectively, show the number of candidate barcodes and the proportion of simulation replicates that were decoded without false discoveries for five selected error-correcting codes as found in simulations of different densities. The Reed-Solomon codes attain the highest robustness by using more images and stronger encoding to reduce the number of ways superimposed barcodes can be recombined into spurious candidate solutions.

Because of the high sensitivity of the FISH-based methods, barcodes can overlap spatially when either many genes are multiplexed or highly expressed genes are probed in an experiment. Previous approaches to solving this density problem have used deterministic super-resolution methods [9–11], expansion microscopy [12, 13], and various image processing optimization models [14–17]. However, none of these methods can resolve ambiguities that arise when multiple barcodes overlap spatially in imperfectly registered data. A previous work suggested a possible graph-based solution to resolve ambiguity from jitter[18]. Here, we introduce a new spots-first decoding method to disentangle overlapping barcodes using global optimization.

After preprocessing the images and fitting dots, we use dynamic programming to identify all sets of aligning dots that could represent a barcode. In dense datasets, recombinations of dots from two or more overlapping barcodes and dots originating from non-specific binding events can give rise to spurious solutions (Fig. 1C). We resolve this ambiguity by using integer-linear programming with the constraint that no two barcodes containing a shared dot are allowed in a solution. The cost of each solution is calculated as a weighted sum of the position variances and log-brightness variances of dots comprising each barcode decoded in the solution and the number of dots in the conflict network that are not decoded in the solution (Fig. 1D).

To measure the robustness of various error-correcting codes to ambiguity arising from spatially overlapping barcodes, we simulated dot locations from multiple overlapping barcodes in a 1-pixel (100*×*100nm) region of interest and evaluated the spots-first decoding results for each error-correcting code (Fig. 1E and Supplementary Fig. 2A). We analyzed seqFISH and MERFISH codes from the literature. SeqFISH used *q*-ary parity check family codes that draw symbols from an alphabet of size *q >* 2 and included one or two parity check symbols [8–10]. MERFISH used binary (*q* = 2) codes: either Hamming codes [2] or non-linear codes [13]. Our simulations showed that the MERFISH and intron seqFISH codes cannot tolerate more than one barcode per region, while the codes used in RNA and DNA seqFISH+ [9, 10] can tolerate two-four spatially overlapping barcodes (Fig. 1E).

To explore whether other codes can provide significant improvements, we analyzed Reed-Solomon (RS) codes [19]. Reed-Solomon codes are also *q*-ary codes. RS codes provide a flexible design framework for designing larger codes with larger minimum distances. The parity check equations in RS codes are designed using abstract algebra structures to ensure that the minimum distance between any two of their codewords (MHD) is one plus the number of parity check symbols, increasing robustness. Simulated data encoded using Reed-Solomon codes can be robustly decoded at densities of greater than five barcodes per search radius (Fig 1E,F). A *q*17 code covering 12,000 genes can achieve decoding of 10 spatially overlapping barcodes with 5% false discovery rate (FDR).

A key determinant of how well an error-correcting code can distinguish many overlapping barcodes in a dense region is how likely dots from overlapping barcodes encoded with that error-correcting code are to recombine into other candidate barcodes. For example, the 2019 MERFISH non-linear code and a subset of the q11n10k7 Reed-Solomon code with 271 weight-four codewords have the same MHD, but the 2019 MERFISH non-linear code produces approximately 20 times as many candidate barcodes in simulations of overlapping barcodes because its codewords are packed more closely in coding space (Fig. 1G). Consistently, simulation replicates for the Reed-Solomon code were much more likely to be decoded without false discoveries than they were for the MERFISH code (Fig. 1H). Specifically, simulated data for 10 overlapping barcodes encoded with the Reed-Solomon code were decoded with an overall FDR of 2%, while simulated data for single barcodes, not spatially overlapping with other barcodes, using the 2019 MERFISH non-linear code were decoded with an FDR of 7% (this was alleviated experimentally by expanding the sample). Reed-Solomon code performance scales well when adding more readout dots per barcode and more readout images. We find that Reed-Solomon codes with larger MHD are more robust to density, but lose robustness when using codewords of weight larger than their MHD and that the additional readout dots increase the number of ways that barcodes can be recombined into spurious solutions (Supplementary Fig. 2).

Even though all codes can be represented in binary form, using symbol blocks with many pseudo-colors (*q*-ary) rather than binary codes increases robustness because only one dot from any given symbol block is allowed in any barcode. This rule reduces the similarity of codewords and reduces ambiguity when barcodes are superimposed. It also means that changing the value of a symbol from one non-zero value to another requires both a removal and an addition of a dot for a change of 1 Hamming distance. Such a change would be measured as 2 Hamming distance in a binary coding scheme. Additionally, thinning codebooks to reduce the number of barcodes under consideration reduces the number of ways to recombine dots into spurious barcodes (Supplementary Fig. 2B).

To validate the simulation results, we applied this decoding method to published seqFISH+ data probing mRNAs of 10,000 genes in NIH 3T3 cells [9]. We measure the sensitivity of our decoding method for a range of parameters by regressing against the average transcript counts of 60 genes measured by sequential single-molecule FISH (255 and 288 cells, respectively) (Fig. 2A,D). Our spots-first processing workflow achieves 46% sensitivity. There was an experiment-wide average of 12 barcodes per micron of cell mask area in each independently encoded optical channel. 27% of decoded barcodes are located within 100 nm of at least one other barcode and would have been discarded by the mutual nearest neighbor method of decoding used in previous seqFISH studies. Another decoding pipeline (Py-FISH) that discards overlaps without attempting to resolve them achieved a sensitivity of 39% with 5% estimated FDR [20], a sensitivity that is lower by almost exactly the proportion of barcodes that are overlapping (27%). We also test an alternative compressed sensing variant of the decoding method on this dataset that uses LASSO regression to choose the subset of candidate barcodes whose expected signals best explain the observed pixel intensities in images of all readout hybridization cycles. It achieves 48% sensitivity at 5% estimated FDR. This method is more robust to spatially overlapping dots in individual images, but more sensitive to registration error than the spots-first approach, precluding its use in the more poorly registered datasets discussed below. Further development of this approach is a promising future direction.

**Fig. 2.**
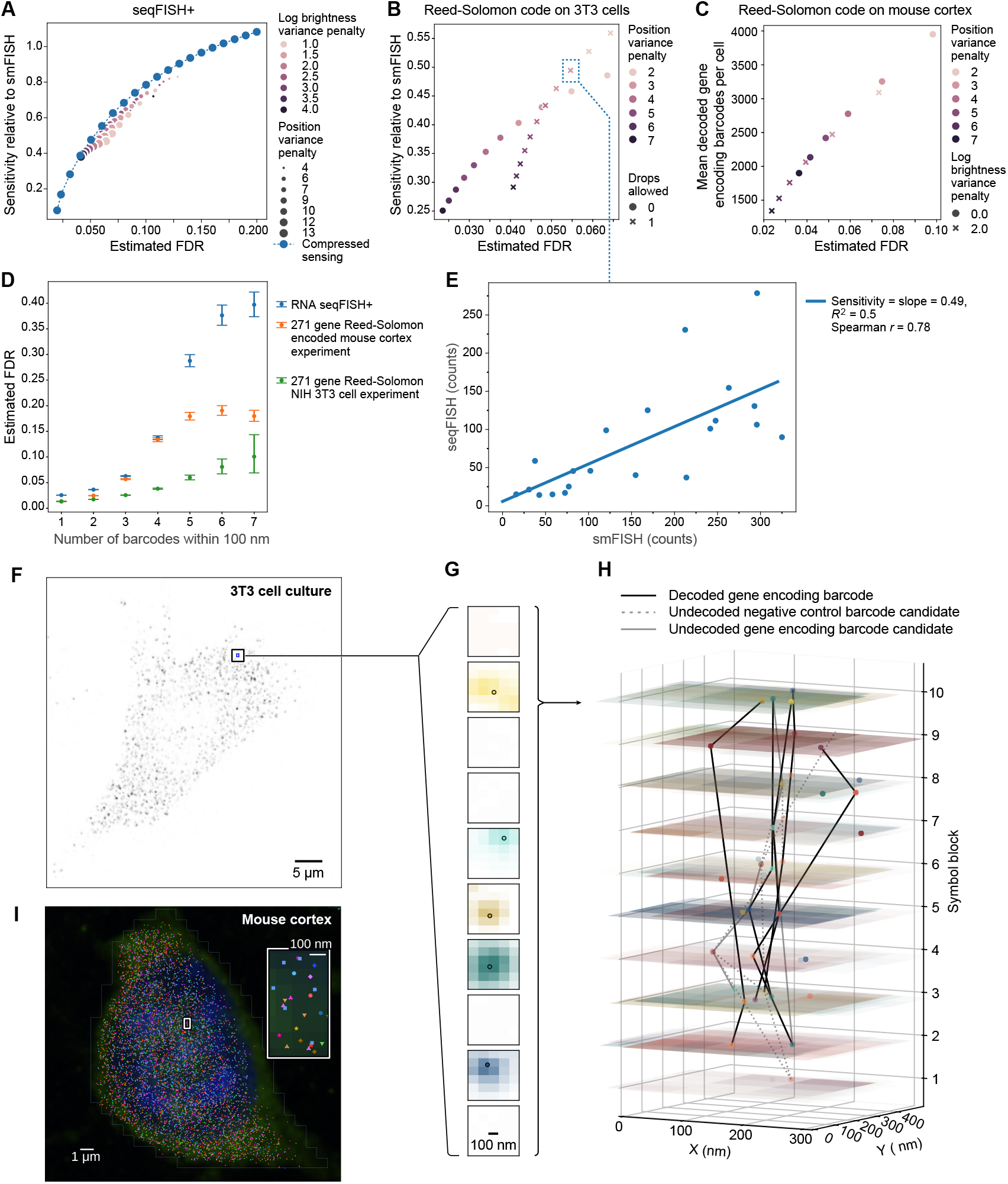
Experimental validation of our untangling approach with RNA seqFISH. A-C) Estimated sensitivity-FDR curves for decoding results on previously published RNA seqFISH+ data in NIH 3T3 fibroblasts (A), a new cross-channel Reed-Solomon encoded experiment in NIH 3T3 fibroblasts (B), and a new single-channel Reed-Solomon encoded mouse cortex experiment probing the same molecules as in the new 3T3 fibroblast experiment (C). Different parameters of the decoding objective functions tune the tradeoff in each experiment. Decoding results for all combinations of parameters in the mouse cortex experiment (C) were evaluated in only nine fields of view to save computation time. D) The estimated FDR of barcodes at different local densities (number of decoded barcodes including self within 100 nm search radius) using position variance penalty of 7 and other penalty parameters of zero for the Reed-Solomon encoded experiments and position variance penalty of 10.5 and log weight variance penalty of 4.0 for the previously published NIH 3T3 cell experiment. Error bars are a 95% confidence interval estimated by non-parametric bootstrap. E) The sensitivity relative to smFISH for the annotated data point in panel B, whose associated statistics were reported in text, was calculated. Decoding sensitivity is measured as the slope of the linear regression fit between average counts of genes found per cell by cross-channel Reed-Solomon encoded seqFISH against the average counts of the same genes found per cell by smFISH on NIH 3T3 cells taken from the same culture. This figure uses barcode counts found using the decoding parameters for which summary statistics are reported in the text. F) A single readout image of a cell from the cross-channel Reed-Solomon encoded 3T3 cell experiment showing dots of the first pseudo-color of the tenth symbol block. G) Images of the ten pseudo-colors of the inset region of the cell in panel F overlaid with fit dot locations in the tenth symbol block which are collapsed into a symbol block image in the top layer of panel H. H) An illustration of decoding results from the inset region of the cell shown in panel F from the cross-channel Reed-Solomon encoded experiment using a codebook including 1829 negative control codewords in addition to the 271 gene encoding codewords. Z-planes represent pseudo-colored symbol block images where each pseudo-color has been registered. Markers denote the fit locations and pseudo-colors of dots in each block. Dots connected by black lines represent decoded gene-encoding barcodes. Dots connected by solid gray lines represent undecoded candidate barcodes encoding for genes. Dots connected by dotted gray lines represent undecoded negative control candidate barcodes. No negative control barcodes were included in the lowest cost decoding solution for this example. I) A maximum projected image of a cell from the mouse cortex experiment with DAPI stain (blue) and poly-T mRNA FISH stain (green) overlaid with locations of decoded barcodes in all z-slices (markers).

To show experimentally that the untangling approach can decode high-density genes, we designed and performed a seqFISH experiment encoded using a Reed-Solomon code. The experiment probed 271 genes expressed at a sum-total of 43,528 FPKM out of a total of 442,744 FPKM expression for all genes in bulk RNA sequencing and an experiment-wide average of approximately 13 barcodes per square micron of cell mask area. Each transcript was read out in 4 out of 100 readout images acquired in 34 hybridization cycles across three optical channels. This contrasts with the previous 10,000 gene seqFISH+ experiment, which consisted of three separately encoded sub-experiments each probing 3,333 or 3,334 genes expressed at a total of ∼41,000 FPKM over 80 hybridization cycles in a single optical channel. Even with chromatic aberration reducing the quality of cross-channel image alignment in the 271 gene experiment, we achieved 49% sensitivity relative to smFISH for 21 selected genes measured in cells taken from the same culture (217 and 218 cells, respectively), detecting a mean of 18,903 transcripts per cell (sample standard deviation=7,572) for all 271 measured genes, with an estimated FDR of 5% (Fig. 2B,D-H and Supplementary Fig. 3). 32% of the barcodes were within 100 nm of another barcode and would have been discarded by mutual nearest neighbors decoding. Per-cell gene correlations calculated using the seqFISH data show a similar structure to that found using smFISH data (Supplementary Fig. 3D-F).

To further validate our method, we used the Reed-Solomon encoded probe set to measure gene expression in the mouse cortex in a single optical channel (Fig. 2C-D,I), finding an average of 2,804 transcripts per cell over all fields of view. A 10X single-cell RNAseq dataset of the mouse cortex in the Allen Brain Atlas found an average of 564 UMI counts per cell for these genes and an average of 11,284 UMI counts per cell for all genes. The tissue was imaged in eight optical z-slices spaced at one micron, and the 2D algorithm was performed on each z-slice. The average barcode density per mask area over all z-slices was six barcodes per square micron using the least stringent decoding parameters in the nine FOVs on which all decoding parameters were tested. We compare the results of our experiment to sample commercial spatial transcriptomics mouse brain datasets acquired using MERFISH (Vizgen)[21], Cosmx (Bruker)[22], and Xenium (10X genomics)[23]. The sample CosMx and Xenium datasets include enough of the same genes as in our experiments to directly compare sensitivity. We find that our method has approximately four and ten times higher sensitivity, respectively (Supplementary Fig. 4A). Furthermore, even though we did not select our gene panel to do so, we were able to predict whether or not each cell in our dataset was a neuron or not with a median confidence of 99.5% using CONCORD label transfer from Allen Brain Cell Atlas single-cell sequencing data (Supplementary Fig. 4B)[24, 25].

While spatial genomics borrows error-correcting codes from digital communications, the two fields operate in different regimes. In telecommunications, errors result from bit-flips in messages, but multiple messages are never sent along the same channel simultaneously. In contrast, the main bottleneck in imaging-based spatial genomics experiments arises from spatially overlapping barcodes and suffers less critically from missing signals, especially with amplification chemistry. The key insight enabling this advance is that error-correcting codes can not only correct transmission errors in telecommunications messages, but also reduce ambiguity arising from spatially overlapping messages in imaging-based spatial genomics data. The finding that spatially overlapping barcodes can be distinguished when they are sufficiently far apart in coding space allows for a new kind of code-based super-resolution microscopy that programs ON and OFF events rather than relying on stochastic switching.

## Author Contributions

J.A.W. and L.C. conceived the idea. J.A.W. wrote the seqFISH data processing packages, seqFISH data processing workflows, simulations, and codebook generation software. L.O. helped with the data processing workflows. J.A.W. and C.L designed the experiments. C.L. performed the experiments and the smFISH data processing. C.L. wrote the experimental methods section and J.A.W. wrote the manuscript with input from all authors.

## Acknowledgements

We thank Jonathan Fox for assistance with animal procedures and tissue collection. We thank Jina Yun for readout probe conjugation and experimental support. We are grateful to Saori Lobbia for maintaining cell cultures and preparing cell samples. We thank Inna-Marie Strazhnik for illustrating our figures. We also thank Victoria Kostina, Matt Thomson, David Van Valen, Mitch Guttman, Linus Eng, and Heather J. Zhou for their helpful comments and feedback. Computations presented here were conducted in the Resnick High Performance Computing Center, a facility supported by Resnick Sustainability Institute at the California Institute of Technology. This work is funded by NIH DP1 NS131408.

## 3 Methods

### 3.1 Fitting dots with ADCG

Dense seqFISH+ images have overlapping dots, which are difficult to distinguish using Laplacian of Gaussians peak finding, the method used in prior works [1, 6, 8]. To better distinguish these dots, we adapted an algorithm that excelled in a benchmarking study for processing dense 2D single-molecule localization microscopy (SMLM) images, Alternating Descent Conditional Gradient (ADCG), to process seqFISH+ image stacks [26, 27]. ADCG fits models of images as linear combinations of Gaussian point spread functions (PSFs) emitted by fluorescent molecules,

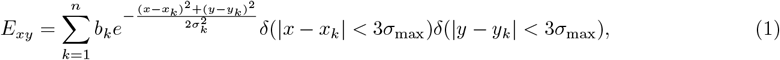

where *E*_*xy*_ denotes the model’s expected intensity of the pixel in the *x*th column and *y*th row in the image; *x*_*k*_, *y*_*k*_, *b*_*k*_, and *σ*_*k*_ are the *x*-coordinate, *y*-coordinate, brightness, and width parameters of the *k*th PSF in the model; *δ* represents an indicator function that saves computation time by not calculating the PSF tails; and *σ*_*min*_ and *σ*_*max*_ bound allowed values of *σ*_*k*_. We include the width as a fit parameter because FISH dots may vary in size due to their distance from the imaging plane and because they are generated by multiple dye molecules attached to an RNA strand of finite size and varying conformation. The fitting procedure uses a least squares loss function

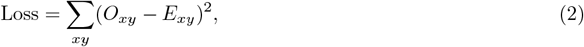

where *O*_*xy*_ denotes the observed intensity of the *xy*th voxel in the image.

ADCG iteratively fits models by adding one new PSF at a time. The first step of each iteration is to approximate the location of the next PSF to add to the model by convolving the image-model residuals with a 2-D Gaussian of width *σ*_min_, then finding the local maxima of the convolved image. Each PSF’s width is approximated by grid search, where linear least squares fits find associated brightness. Second, gradient descents refine the coordinate, then the width and brightness estimates. We do not differentiate the indicator functions when calculating gradients of the loss, but we retain them in our gradient function to save calculation time. Third, repeated gradient descents on the coordinates then brightness and width of all PSFs adjusts the parameters of all PSFs in the model until the loss converges or for a maximum number of iterations. Finally, PSFs with brightness below the minimum allowed are removed. PSFs are added to the model until no new PSFs with brightness larger than the minimum allowed are found, the improvement in the objective function is below a threshold, or a maximum allowed number of PSFs have been added to the model.

When the algorithm terminates because the brightness or objective change is below a threshold, it discards the changes from the final iteration to avoid overfitting. For computational efficiency, we break up images into overlapping tiles, run the fit on each tile, and then piece the results from each tile together for the full image, removing duplicate PSFs found in the overlapping regions. Our implementation of this algorithm is available at https://github.com/CaiGroup/SeqFISH_ADCG.jl.

### 3.2 Fiducial marker matching registration

Dots found in different symbol blocks of seqFISH experiments must be aligned to correct for drift over the course of the experiment and recognize which were read out from the same barcode. Previous studies [1, 2, 8, 28] have aligned hybridization images using phase cross-correlation [29] on independent images of either DAPI stains or fiducial markers acquired concurrently but at a different wavelength than the readout images. This approach introduces errors from optical aberration, differences in light path alignment, and movement of the microscope objective between scans. We reduce these sources of error by aligning using diffraction-limited objects as fiducial markers in the readout images identified with a pattern matching algorithm [10]. We then register images in the same optical channel as each other using translations and images in different optical channels using affine transformations.

Fiducial markers do not move relative to the slide, so they appear in a constellation, i.e. a fixed pattern, that drifts with the field of view in the readout images (Supplementary Fig 5A-C). Our algorithm searches for the dots in a fiducial marker constellation found in a reference image of only the fiducial markers among all dots found in readout images. The constellation of fiducial markers is identified by matching the relative position vectors between pairs of fiducial markers. Each matching pair of separation vectors in the reference and readout images votes to classify the dots at their ends as matches. This is similar to another pattern matching algorithm that matches triangles, rather than vectors [30].

#### Algorithm 1 Matching fiducial markers algorithm definitions

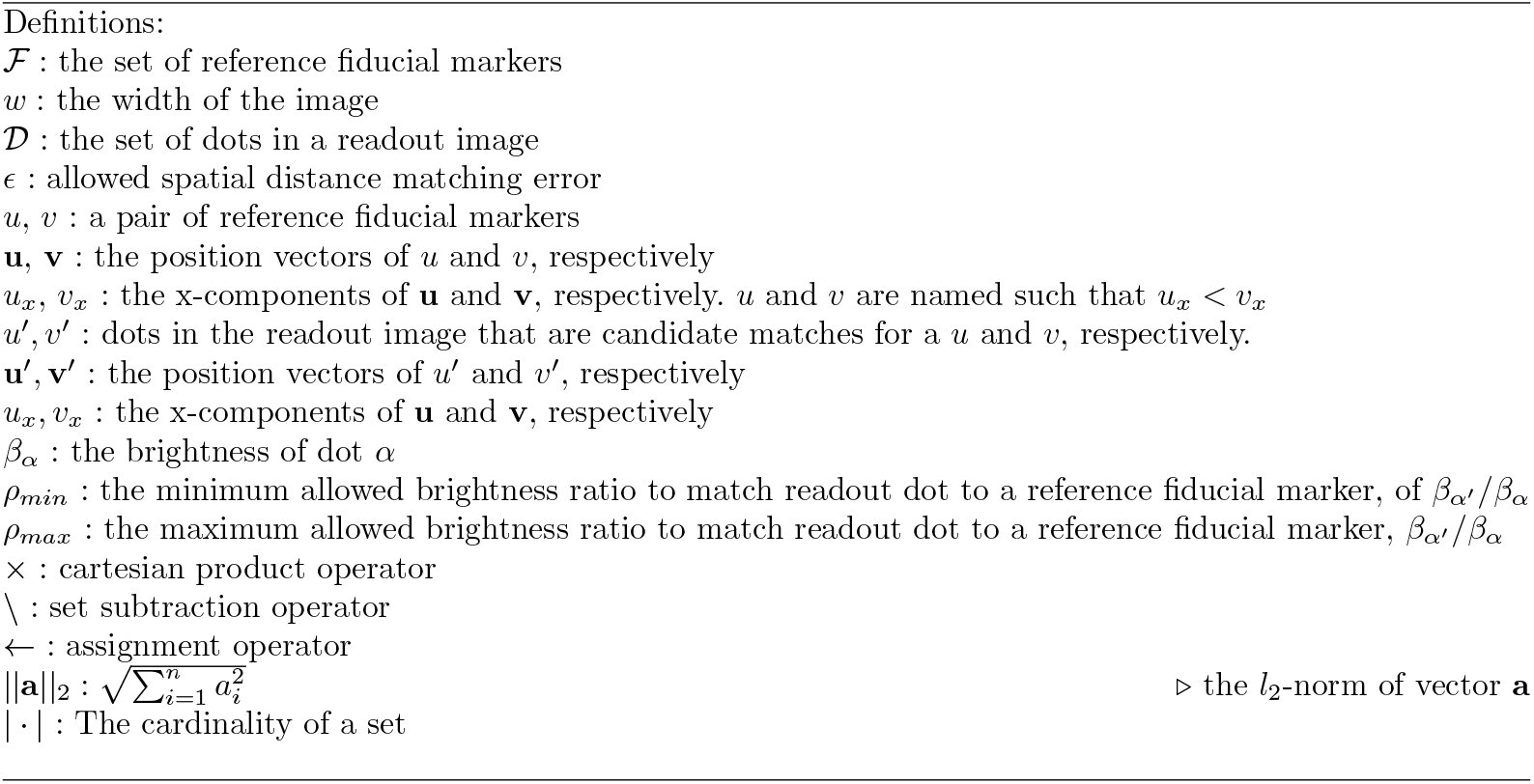

#### Algorithm 1 Fiducial marker matching algorithm

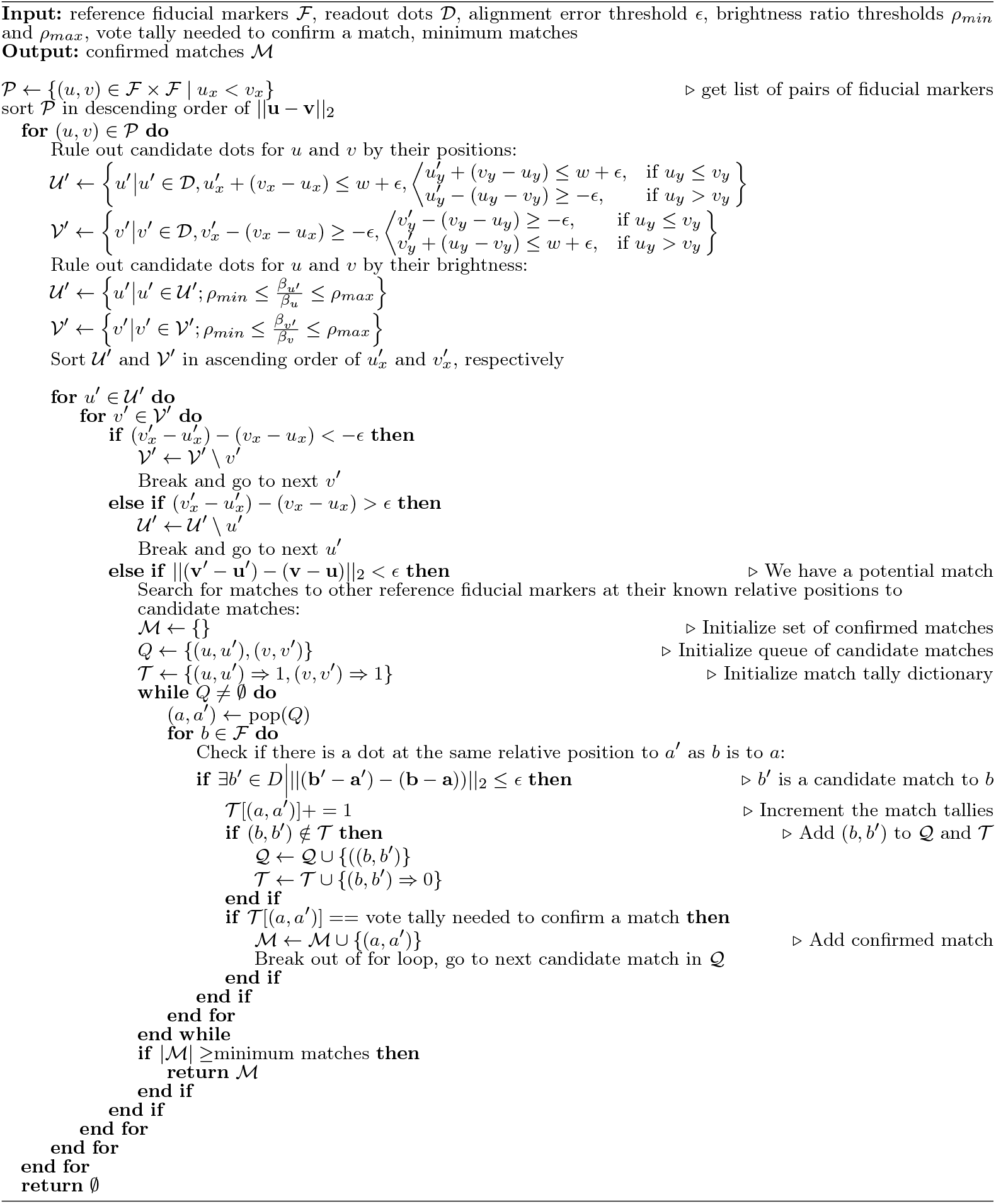

The most challenging task is to find the first matching pair of separation vectors. We begin by assembling a list of the most distant pairs of fiducial markers from the reference image, excluding any fiducial marker that is within the allowed matching error distance, *ϵ*, of another fiducial marker. We denote the two fiducial markers in a pair as *u* and *v* such that the *x*-component, *u*_*x*_, of *u*’s position vector, **u**, is less than the *x*-component, *u*_*x*_, of υ’s position vector, **v**. The list is sorted in descending order of ||**v − u**||_2_. Initially searching for the most distant pairs of fiducial markers lets us narrow the region of the readout image where we must search for *u* and υ to around the corners.

For each (*u, υ*) pair, we rule out dots as candidates for *u* and υ. We rule out *u*′ and υ′ candidates at locations in the image where their partner dot would be off the image. We rule out more dots by setting bounds on the brightness ratio between fiducial markers in the reference image and readout image. When drifts are small, we can also restrict the search to dots within a maximum allowed shift between images. To find an initial match, we sort the lists of dots in the readout image that may match to *u* and υ by their *x*-coordinate from lowest to highest. We denote individual candidate dots *u*′ and υ′. We first compare the *x*-components of the position vectors separating each *u*′ and υ′, **v**′ − **u**′, with the position vector separating reference dots, **v** − **u**. We start with the first *u*′ and υ′ candidates in the lists, then proceed through the following logical flow: if 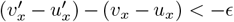, then υ′ cannot match υ, so we discard it and start again at the beginning of the revised lists. Likewise, if 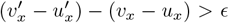, then *u*′ cannot match *u*, so we discard it and start at the beginning of the revised lists. If ||(**v**′ − **u**′) − (**v** − **u**) ||_2_ *> ϵ*, then we check the next υ′, in the list. If|| (**v**′ − **u**′) (**v** − **u**) ||_2_ *< ϵ*, the relative position vectors match and we proceed to search for the rest of the fiducial markers relative to it. When there is fewer than one candidate dot for every error window of 2*ϵ*, this algorithm will find matches in *O*(𝓃) time.

To begin the search for the rest of the fiducial markers in the readout image, we initialize a queue containing the first two candidate fiducial marker matches, (*u, u*′) and (υ, υ′). We then begin taking fiducial marker matched dots from the queue and searching for other fiducial markers from the reference pattern at their known relative position from the already matched dot in the readout image with a KDTree. For example, if we already have matched the dot *a*′ to the fiducial marker *a*, we search for another dot *b*′ that matches another fiducial marker *b* such that || (**b**′ **a**′) − (**b a**)) ||_2_ *< ϵ*. If such a *b*′ is found, the candidate-reference pair, (*b, b*′), is added to the candidate match queue. Each (*b, b*′) candidate match that is found relative to an (*a, a*′) candidate match is a matching edge that votes for the (*a, a*′) and (*b, b*′) matches. Once an (*a, a*′) candidate matching pair has received ten votes from matching separation vectors, the match is confirmed, and then the algorithm proceeds to search for matches with the next (*a, a*′) candidate-reference pair in the queue. If no (*b, b*′) candidate matches are found in the first ten searches from a given (*a, a*′) pair of candidate matches, the (*a, a*′) candidate match is discarded. If fewer than five matches have been confirmed with ten matching edge votes after exhaustively searching from initial (*u, u*′) and (υ, υ′) candidate matches, all matches are discarded and the algorithm searches for new initial matches.

Search parameters may either be set manually by the user or the algorithm may first conduct a grid search for parameters to match the fiducial markers in two reference images of only the fiducial markers acquired at the beginning and end of the experiment. In the latter case, the algorithm chooses the strictest value for the search radius in which it finds 90% of the matches that if found with the least strict parameters, the minimum brightness considered for a match to an initial dot (*ρ*_*min*_) to be 50% less than the ratio of the greatest reduction in brightness from the initial to the final reference image, and the maximum allowed brightness ratio of matches to initial fiducial markers (*ρ*_*max*_) to be the maximum observed brightness ratio of initial to final reference dot. Our implementation of this algorithm is available at https://github.com/CaiGroup/seqfish_fm_match.

### 3.4 SeqFISH representation of error-correcting codes

Reed-Solomon codes are a family of error-correcting codes with an elegant mathematical construction that are maximum distance separable: the minimum Hamming distance between any two codewords (MHD) in the error-correcting code is the maximum possible for a linear error-correcting code given the number of parity check symbols it contains. To represent Reed-Solomon codes using seqFISH, we use only codewords with a fixed weight (number of non-zero symbols) and probe only non-zero symbols in each codeword. Zero-valued symbols are represented by symbol blocks without a dot. We encoded our experiment using weight-four codewords from the Reed-Solomon code with 10 symbols (*n* = 10), including seven information symbols (*k* = 7) and three parity check symbols (*n* − *k* = 3), from an alphabet of size 11 (*q* = 11). Our simulations include codewords from other Reed-Solomon codes using codewords of weight four, five, and six (Fig. 1E and Supplementary Fig. 2). Our supplementary simulations include a Reed-Solomon code whose symbols are elements of the extended finite field of eight elements. For this and other Reed-Solomon codes defined over extended finite fields, the symbol value represented by a dot is the primitive element of the finite field exponentiated by the dot’s pseudo-color number minus one.

### 3.4 Generating codebooks and parity check matrices

Reed-Solomon codes are defined using the abstract algebra structures of polynomial rings over finite fields. An equivalent representation to the generator polynomial is a generator matrix with elements from the same finite field as the generator polynomial coefficients [31]. To compute the codewords with the desired number of non-zero symbols, we find the systematic form (row reduced echelon form) of the generator matrix, which computes codewords that have parity check symbols appended to the information symbols in the message.

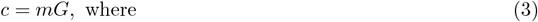

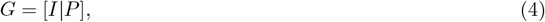

*m* is a message vector containing the information symbols encoding a gene, *c* is the codeword that consists of *m* with parity check symbols appended. *G* is the generator matrix in systematic form, which consists of an identity matrix concatenated with columns representing the parity check equations. Using the systematic form of the generator matrix allows us to consider only messages with the desired number of non-zero information symbols or fewer, compute the parity check symbols for the message, then discard codewords that have more than the desired number of non-zero symbols. We then check that we found the same number of codewords of the desired weight as predicted by the theoretical weight enumeration formula [32]. We provide notebooks implementing Reed-Solomon codeword generation on both CPUs and GPUs for various kinds of Reed-Solomon codes https://github.com/CaiGroup/UntanglingBarcodes/tree/main/codebook_generation/get_RS_codebooks.

### 3.5 Assigning genes to codewords

We optimize the assignment of genes to codewords to minimize the maximum sum FPKM expression of all genes probed in any single hybridization image. Alignment across optical channels is worse than alignment within a single optical channel, so after assigning genes to codewords, we assigned symbol block and pseudocolor values to hybridization and optical channel images to maximize the number of barcodes that are probed in the same optical channel either three out of four or four out of four times. Notebooks are available at https://github.com/CaiGroup/UntanglingBarcodes/tree/main/codebookgeneration/assign_genes_to_codewords.

### 3.6 Identifying candidate barcodes with syndrome decoding

To identify candidate barcodes, we use dynamic programming to find dots that align in different symbol blocks whose pseudo-color values satisfy the error-correcting code’s parity check equations. First, we illustrate the algorithm using the error-correcting code from the original seqFISH+ experiment, a *q*-ary parity check code with four symbol blocks and 20 pseudo-colors [9]. All combinations of pseudo-colors are allowed in the first three symbol blocks, but the final pseudo-color is determined by the parity check equation:

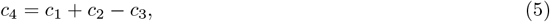

where, *c*_*i*_ denotes the value of the *i*th symbol in an error-correcting code. When describing a seqFISH experiment that transmits information using this code, we say that *c*_*i*_ denotes the pseudo-color number of an aligning dot in symbol block *i*. Addition and subtraction operations are modulo 20, the number of pseudocolors. With this code, any three aligning dots in each of the first three symbol blocks with any combination of pseudo-color values may represent a barcode. In the terminology of error-correcting codes, these dots represent information symbols that alone are sufficient to represent all the information encoded in a message. Dots in the final symbol block represent parity check symbols, which add robustness to the message. The parity check equation defines the syndrome,

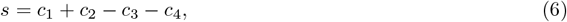

such that the syndrome, *s*, will be 0 if and only if the sequence of symbols represented by the aligning dots’ pseudo-colors is a codeword. In general, linear error-correcting codes are defined by a parity check matrix, *H*, such that

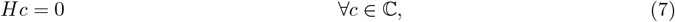

where ℂ is the set of codewords in the code. Codewords, *c*, are sequences of symbols drawn from an alphabet of size *q*. Arithmetic operations are modulo-*q*. The parity check code used in reference [9] uses a parity check matrix,

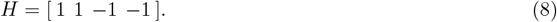

The parity check equation defines the syndrome,

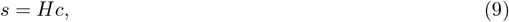

such that all elements of *s* will be 0 if and only if the sequence of symbols represented by the aligning dots is a codeword. The linearity of the syndrome computation allows all candidate barcodes to be found with dynamic programming, a technique that is also known by the more descriptive name, recursive optimization. This procedure involves breaking the matrix multiplication summing operations into steps and saving intermediate sums to reduce the number of arithmetic operations that must be performed.

We take a divide and conquer approach to efficiently compute syndromes and identify candidate barcodes through independent searches in overlapping square tile regions covering the image. For a collection of dots to qualify as a valid candidate barcode, all its dots must align to within the search radius of one another. To guarantee that all valid candidate barcodes are found, a circle with diameter equal to the search radius must be guaranteed to be contained entirely in at least one tile of the tile cover of the image. We achieve this guarantee by defining a tile cover of the image composed of square tiles of width twice the search radius centered at each point in a square grid with spacing of one search radius. This leads to some individual barcodes being found in multiple tiles, but reduces overall computational cost by eliminating chains of subsequently aligning dots that extend long distances and reducing the number of intermediate sums that need to be held in memory concurrently. After finding candidate barcodes in each tile, we remove duplicate candidate barcodes found in multiple tiles.

We now use a recursive framework to describe the dynamic programming algorithm performed on dots in each tile region for the simpler case where barcodes are probed in every symbol block (Supplementary Fig. 1A). In the initialization step, we assign a partial syndrome sum array to each dot in the first symbol block that contains the pseudo-color number (symbol value) of that dot as its lone element. We then proceed to construct partial syndrome sum arrays for dots in each subsequent symbol block. To construct the partial syndrome sum array for each dot, we first use a KDTree search to find each dot’s neighboring dots in the previous symbol block that align to within a search radius. The partial syndrome sum arrays assigned to the aligning dots are concatenated. Finally, the searching dot’s pseudo-color is added element-wise (or subtracted) from the concatenated array, resulting in that dot’s partial syndrome sum array. Syndromes with value 0 in the final symbol block indicate that a chain of subsequently aligning dots has pseudo-colors that are consistent with a valid barcode. We can trace the dots that contributed to zero-valued syndromes. Traced collections of dots for which all dots are within the search radius of each other are valid candidate barcodes. Traced collections of dots for which any two dots are farther apart from one another than the search radius are discarded.

#### Algorithm 2 Syndrome decoding for single parity check codes: Definitions

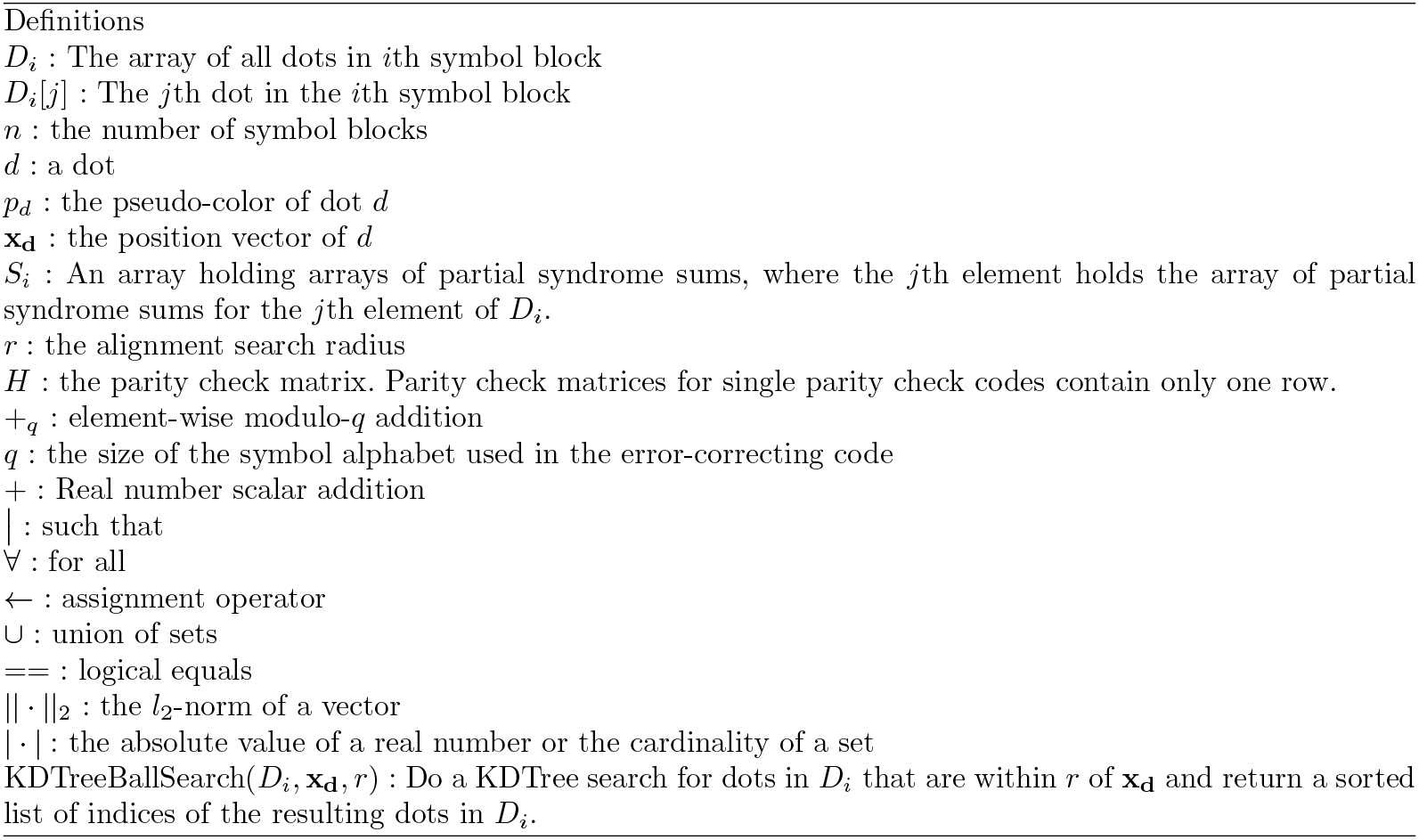

#### Algorithm 2 Syndrome decoding for single parity check codes by recursive sums

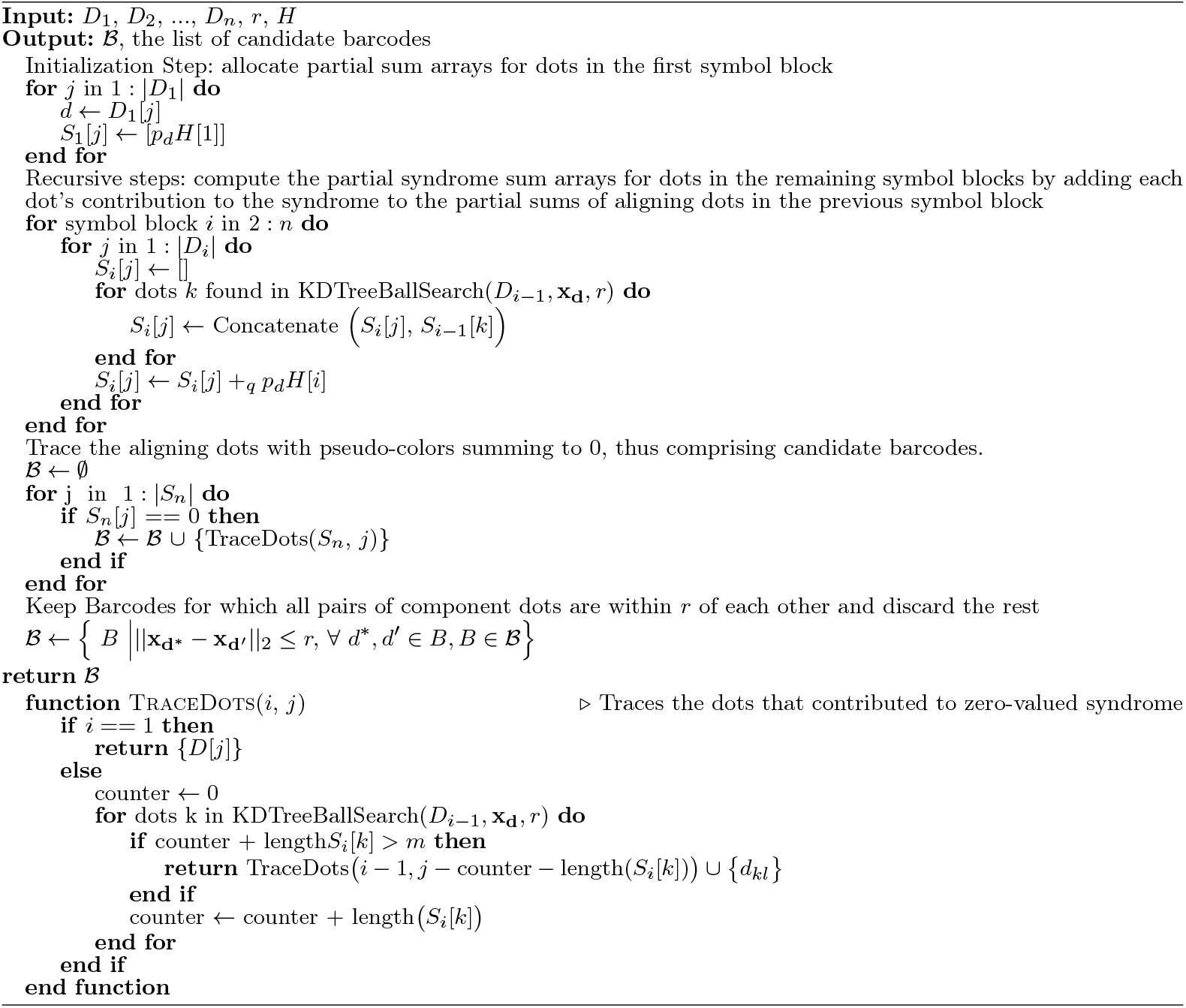

Now we describe the procedure for the more general and complex case where barcodes are not probed in every symbol block (Supplementary Fig. 1B). In this case, we must align dots from each symbol block with dots from multiple previous symbol blocks and track the number of dots contributing to each partial sum. To do this, partial sums for each dot are stored in one of *w* arrays, where *w* is the weight of the codewords used in the experiment, i.e., the number of dots readout from each barcode. Elements in the *i*th of the *w* arrays hold partial sums of pseudo-colors from *i* subsequently aligning dots in different earlier symbol blocks. The first array of partial sums for each dot contains only that dot’s contribution to a syndrome. The *i*th partial sums array for each dot, where *i >* 1, is found by concatenating the (*i* − 1)th nested syndrome partial sum array for dots in previous symbol blocks within the search radius, then adding the syndrome contribution of the searching dot element-wise. The *w*th array holds the syndromes for paths of aligning dots. Entries in the *w*th array with value 0 represent candidate barcodes and we can trace the dots that contributed to it. When allowing incomplete barcodes with 1 missing dot, all paths of length *w*− 1 are traced and searched in a BKTree of codewords to check whether they are Hamming distance 1 from a valid codeword. Our implementation of this algorithm is available at https://github.com/CaiGroup/SeqFISHSyndromeDecoding.jl.

### 3.7 Resolving conflicting decoding solutions with integer programming

In dense experiments, syndrome decoding will yield conflicting barcode candidates that include at least one of the same dots (Fig. 1C). The densest regions may have large networks of conflicting barcode candidates. To find the non-conflicting barcodes that best explain the observed dots (Fig. 1D), we define a cost function for each feasible solution

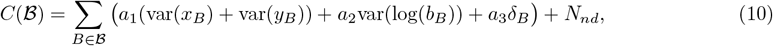

where ℬ is the set of all barcode candidates in the network; *B* is a single barcode candidate; *x*_*B*_, *y*_*B*_, and *b*_*B*_ are the sets of *x*-coordinates, *y*-coordinates, and brightness of each dot in the candidate barcode *B*; *δ*_*B*_ is an indicator function that is equal to 0 if the barcode *B* is complete or 1 if the barcode is missing a dot; and *N*_*nd*_ is the number of dots in the network that are not in a barcode candidate chosen by the solution. var is the sample variance of its input set defined as

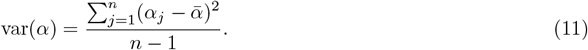

The logarithm on the set of brightnesses of the dots in a barcode is applied element-wise. *a*_1_, the lateral position variance penalty; *a*_2_, the brightness variance penalty; and *a*_3_, the missing dot penalty, are user set parameters. We use an off-the-shelf integer programming solver to find the lowest cost, non-conflicting set of barcode candidates [33] (Gurobi). We solve the optimization problem as written in standard form:

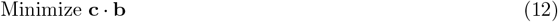

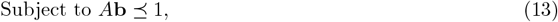

where **b** is a vector of indicator variables for each candidate barcode, **c** is the vector of corresponding costs for each candidate barcode where the cost of the barcode of index *i* is calculated as

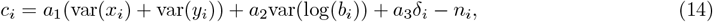

where *n*_*i*_ is the number of dots in the *i*th barcode, ⪯ is the element-wise less than or equal to comparator, and *A* is the adjacency matrix for the candidate barcodes. *x*_*i*_, *y*_*i*_, *b*_*i*_, *δ*_*i*_, *a*_1_, *a*_2_, and *a*_3_ are defined as in equation 10. Barcodes that are composed of at least one of the same dots are neighbors in *A*. Barcode candidates with non-negative costs according to equation 14 are discarded. We implemented a function to perform this optimization in our package located at https://github.com/CaiGroup/SeqFISHSyndromeDecoding.jl.

To apply this to experimental data, we set the search radius to four pixels in our 3T3 cell experiment spots-first workflows and three pixels in the mouse cortex spots-first workflow. We discard candidate barcode conflict networks that have more than 2500 candidate barcodes and have a ratio of candidate barcodes to bounding box area greater than 10 for the seqFISH+ reanalysis or more than 700 candidate barcodes and a ratio of candidate barcodes to bounding box area greater than 700 for the cross-channel Reed-Solomon encoded 3T3 cell experiment and mouse cortex experiments. We tried a range of reasonable values for the cost parameters to gauge the trade-off between sensitivity and false discovery rate (FDR) and plotted ROC-like sensitivity-FDR curves (Fig. 2A-C).

### 3.8 SeqFISH+ 2019 spots-first image processing workflow

We use the snakemake workflow manager [34] to define a reproducible workflow for processing seqFISH images. The seqFISH+ 2019 experimental data is divided into three independently coded optical channels each encoding 3,333 or 3,334 genes in 80 hybridization images divided into four symbol blocks of 20 pseudocolors. This experimental design avoided error from cross-channel image registration and reduced ambiguity by partitioning barcodes that may have overlapped into independent subdivisions. The workflow first fits fluorescent beads used as fiducial markers with ADCG in all images, then finds the translations of readout images relative to a bead-only reference image of the same optical channel by matching fiducial marker patterns. Before fitting dimmer FISH dots, the workflow removes the fiducial markers and autofluorescence from readout images by pixel-wise subtracting the aligned reference images from them. It then subtracts the scattering background intensity estimated by applying a median filter with a 5 *×* 5 pixel kernel and then using the rolling ball algorithm with a radius of 3.3 pixels [35] and masks out intensity from pixels between cells. The same hand-drawn masks used in the original study are used [9]. The workflow then finds FISH dots in the preprocessed images with ADCG, aligns their fit coordinates, splits them into groups aligning to each cell mask, and then uses syndrome decoding to infer which barcodes generated the PSFs in each cell. Only complete barcodes are allowed. The implementation of this workflow is available at https://github.com/CaiGroup/UntanglingBarcodes/tree/main/real_data_processing_workflows/2019_seqfishplus_reprocessing.

### 3.9 Cross-channel Reed-Solomon encoded seqFISH workflow

Our Reed-Solomon encoded experiment uses 100 image stacks acquired in three optical channels and 34 hybridization cycles to represent ten symbol blocks of ten pseudo-colors. 2D segmentations were drawn manually with Cellpose[36] from FISH images of cells hybridized with poly-T probes. First, beads used as fiducial markers were fit in the brightest *z*-slice (4th *z*-slice) where cells were masked out by segmentation masks. Beads in reference images containing only the beads and no readout probes were matched between optical channels and to beads found in images with readouts in the same optical channel. The optical channel with the most fiducial markers from the readout images matched to corresponding fiducial markers in its reference image was used as the reference optical channel for each position. Images in the reference optical channel were registered to their reference image by translation, and images in the other optical channels were registered by affine transformation to the reference optical channel image. Background subtraction for images in each optical channel was performed similarly to the 2019 seqFISH+ experiment processing workflow, but in 3D. After background subtraction, readout dots were fit using 2D ADCG in the brightest (4th) *z*-slice. For this experiment, the increased minimum distance of the code allows incomplete barcodes with only three dots (one missing dot) and the missing dot penalty (*a*_3_ in equation 10 and 14) is set to 2. The implementation of this workflow is available at https://github.com/CaiGroup/UntanglingBarcodes/tree/main/real_data_processing_workflows/Reed-Solomon_encoded_experiment_workflow.

### 3.10 Mouse cortex processing workflow

The mouse experiment used the same genes and codebook as in the cross-channel Reed-Solomon encoded 3T3 cell experiment, however all images were acquired in 561 optical channel using Cy3B fluorescent dye. Z-planes were acquired at intervals of 1 micron. Fiducial marker matching was done in the sixth z-slice to calculate translational offsets. Segmentations were computed on 1/8th downsampled image stacks of the cells with DAPI staining and Poly-T FISH staining using Cellpose-SAM [37]. 2D spots-first decoding was performed on each z-plane independently using a search radius of three pixels. The sensitivity-estimated FDR curve was calculated using the first nine fields of view. The remaining fields of view were decoded with lateral variance penalty of five and log brightness variance penalty of zero. The implementation of this workflow is available at https://github.com/CaiGroup/UntanglingBarcodes/tree/main/real_data_processing_workflows/mouse_cortex

### 3.11 Label transfer in mouse cortex

Cell types in seqFISH mouse cortex data were assigned using CONCORD to transfer labels from Allen Brain Atlas 10X single-cell sequencing data (Supplemental Fig. 4B). Cells in the 10X dataset with fewer than 10 genes, with missing class labels, or that were members of a cell class with fewer than nine cells of that class were removed. Expression counts were normalized in the remaining cells. Cells in the seqFISH dataset with fewer than 100 genes or that were in the manually annotated PIA layer region were discarded. PIA layer cells are difficult to dissociate in single-cell sequencing protocols and are thus likely underrepresented in 10X datasets, leading to poor label transfer results for these cells. Since some cell types are far more represented in the 10X data than others, the training dataset was balanced by choosing a random subsample of 1000 cells for each cell type class, of which there were more than 1000 cells. We trained ten models, each with different balanced subsamples of the training dataset, using only the genes that were profiled in our seqFISH experiment. We used each CONCORD model to predict the cell types of each cell in the seqFISH dataset. We aggregate classes of neurons into two superclasses of GABAergic neurons and glutamatergic neurons. We then find cells that have consensus cell type predictions from all models.

### 3.12 Compressed sensing decoding

The spots-first decoding procedure described above cannot distinguish spatially overlapping dots generated from two or more transcripts in the same optical image. To address this, we develop an alternate decoder that uses a compressed sensing framework to fit barcodes in all optical images simultaneously. The drawbacks compared to the spots-first approach are that candidate dots must be located on a predefined grid and that the computational complexity increases more quickly with increasing search radius, which limits its capacity to handle larger readout dot jitter or registration error between readout images. These limitations may be overcome with future work.

After performing image registration and preprocessing as described for the spots-first workflows, the next step is to identify candidate dot locations in each readout image. Preliminary candidate dot locations, 𝒟, are found by convolving (i.e., blurring) readout images with a Gaussian kernel (the point spread function). Centers of pixels in the convolved (blurred) image with value greater than a threshold are the preliminary candidate dot coordinates,

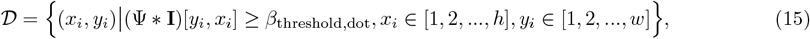

where **I** is the image matrix of dimension *h × w*,

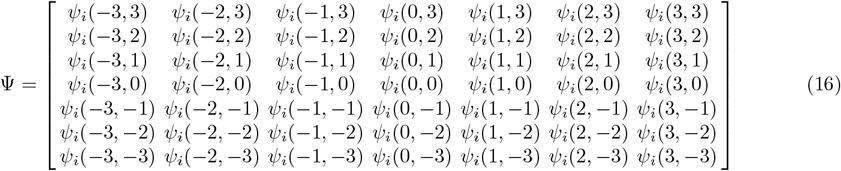

is the Gaussian kernel, ∗ is the convolution operator, (Ψ ∗ **I**)[*y*_*i*_, *x*_*i*_] is the value of the pixel in the *y*_*i*_th row and *x*_*i*_th column of the convolved image, and *β*_threshold,dot_ is a user defined threshold. We use a 2D Gaussian point spread function defined as

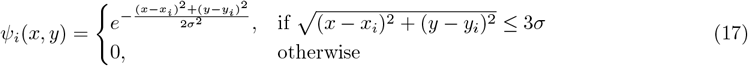

where *x*_*i*_ are *y*_*i*_ are the *x*-coordinate and *y*-coordinate of the center of the point spread function and *σ* is the width parameter.

Candidate locations identified by thresholding the convoluted image are further thinned using a LASSO regression model similar to a model that has previously been used for single-molecule localization microscopy [38]. Column vectors, **d**_**i**_, of the sensing matrix, *A*, are flattened representations of the signal expected to be readout from a molecule centered at coordinates (*x*_*i*_, *y*_*i*_) ∈ 𝒟. We can write the optimization problem as

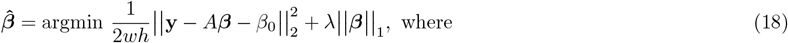

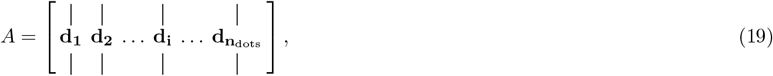

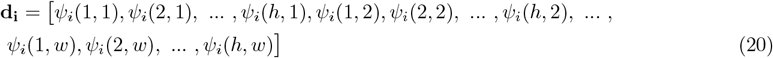

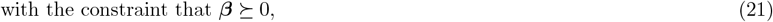

where **y** is the vector of observed pixel values in the image, ***β*** is the coefficient vector, *β*_0_ is the (scalar) intercept, *λ* is the regularization parameter, *n*_dots_ is the number of candidate dots, *h* is the number of rows, *w* is the number of columns of pixels in the region of interest, and ⪰ is the element-wise greater than or equal to symbol. To reduce computational cost, rows of *A* with all zeros and the corresponding elements of **y** and ***β*** are removed. Only candidate dots with *β*_*i*_ ≥ *β*_threshold,dot_ at the smallest value of *λ* for which *β*_0_ is non-negative are kept. Remaining candidate dot locations are then registered using translations calculated from matched fiducial markers, and candidate barcodes are identified using the syndrome decoding algorithm as described for the spots-first workflow with a user-supplied search radius of *r*. This yields a list of candidate barcodes

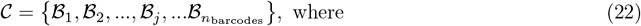

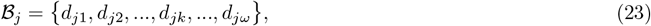

where ℬ_*j*_ denotes the *j*th candidate barcode, *d*_*jk*_ denotes the *k*th candidate dot in the *j*th candidate barcode, and *ω* is the weight of the codewords, i.e. the number of dots in a barcode.

Next, for each cell, a LASSO model is constructed to select candidate barcodes. In this model, the **y** vector contains the pixel values of all readout images of the cell and columns of the sensing matrix, *A*, indicate the expected illumination patterns for each of the *n*_barcodes_ candidate barcodes in all images

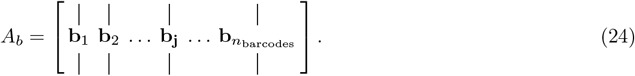

The column vectors, **b**_**j**_, representing the signal expected to be emitted by the *j*th candidate barcode in all images, are constructed by concatenating a sequence of vectors representing the expected signal, **s**_**i**_, from the candidate barcode in each image,

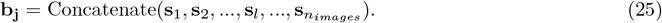

The expected signal for images is

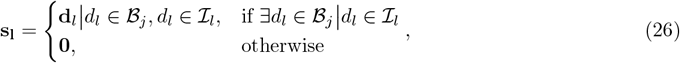

where **d**_**l**_ is the flattened dot signal vector defined in equation 20 for the component dot, *d*_*l*_, in the barcode, ℬ_*j*_, and in ℐ_*l*_, the set of dots in the *l*th image, **0** is a vector of zeros of length *wh*, the number of pixels in each image. The LASSO optimization problem for selecting barcodes is

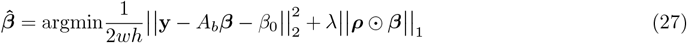

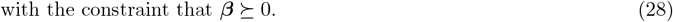

The ⊙ operator represents element-wise multiplication, and ⪰ is the element-wise greater than or equal to symbol. The LASSO penalty for each barcode is modulated by the variance of the positions of the candidate dots comprising each barcode. ***ρ*** is a vector of penalties for each barcode and its *j*th element is calculated as

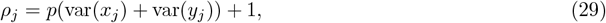

where var is defined in equation 11, and *x*_*j*_ and *y*_*j*_ are the registered *x* and *y*-coordinates of the dots in the *j*th candidate barcode respectively, and *p* is a parameter controlling the severity of the penalty. Again, to reduce computational cost, we remove rows of *A* that are all zeros and corresponding elements of **y** and ***β***. The glmnet package returns a LASSO path with many values of the LASSO penalty, *λ*, starting with the largest value that returns a non-null model, *λ*_0_, and ending with *λ* = *λ*_0_*/*1000 [39]. The LASSO path includes a *B* matrix where each *i*th column vector is the fit value of 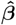for the *i*th value of *λ*.

Compressed sensing relies on the sensing matrix having low mutual coherence (i.e., distinct columns) to guarantee unique and accurate solutions [40, 41]. However, matrices *A*_*b*_ as defined in equation 24 will have many highly coherent columns that represent slight variations in candidate dot locations to explain the signal emitted from an individual molecule. Many of these overlapping candidate barcodes for transcripts of the same gene will be selected at lower values of *λ*, diluting their coefficient values. Thus, at any particular value of *λ*, candidate barcode *j* that have had a fit coefficient value greater than the threshold, 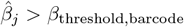, at any *λ* ≥ *λ*_*m*_ are considered selected at *λ*_*m*_. We compensate for this by removing duplicate overlapping barcodes for the same gene selected at each value of *λ*. To identify duplicate barcodes, we first construct networks of selected barcodes that encode each kind of RNA molecule. Barcodes in a network are neighbors if they are centered within the decoding search radius, *r*, of each other. Of all barcodes in a connected network, we only keep the barcode that is included in the model with the largest fit coefficient value, *β*_*j*_, at the largest *λ* where any of the barcodes in the network are included in the model. The other selected barcodes in the network of barcodes encoding the same gene are discarded. In future implementations, it may be desirable to remove duplicate overlapping barcodes either using *l*_0_-regulated regression as demonstrated in supplementary simulations (Supplementary Fig. 6B, Supplementary Note 3) or to group overlapping barcodes in a grouped LASSO model to learn the spot shape of non-diffraction-limited target molecules [42–44].

The sensitivity-estimated FDR curve shown in Fig. 2A is calculated by first running the above algorithm with every combination of candidate *β*_threshold,dot_, *r, ρ*, and *β*_threshold,barcode_ requested by the user. Then the best estimated FDR curve for each cell is found by keeping the results only for which there are no other parameter sets with a lower estimated FDR and a higher sensitivity. The overall estimated FDR-sensitivity curve is found by averaging the decoding results from each cell when decoded with the value of the parameters that yielded the closest estimated FDR to the target value. The implementation of this workflow is available at https://github.com/CaiGroup/UntanglingBarcodes/tree/main/real_data_processing_workflows/compressed_sensing

### 3.13 Simulations to determine density robustness of codes

Imageless simulations of decoding performance with various error-correcting codes included seqFISH *q*-ary parity check codes, the Hamming code used in the original MERFISH experiment [2], the non-linear sequence space cover code used in later MERFISH experiments [13], Reed-Solomon Codes, and a new non-linear code (Fig. 1E, Supplementary Fig 2, Supplementary Note 1). The units of all distances in this section are pixels, which in our experiments are approximately 100 nm in width. Simulation workflows first generate the ground truth (the coordinates and identity of each simulated transcript) for each replicate of each condition for each error-correcting code. *x* and *y*-coordinates of each transcript were drawn from a uniform distribution from 0 to 1. The codeword of each object was randomly selected from the given codebook with equal probability for each codeword. Then, the workflow generated simulated observed readout dot locations from each simulated transcript within the appropriate symbol block and of the appropriate pseudo-color. The localization error of each simulated readout dot from the simulated true location of each simulated transcript was drawn from a 2D normal distribution with *σ* = 0.5. Each readout dot had a 1/20 probability of being dropped in the simulation data. Dots of each pseudo-color of each symbol block have a 1/50 probability of including a non-specific dot with *x* and *y*-coordinates drawn from a uniform distribution from 0 to 1. Simulated dots were fed into the SeqFISHSyndromeDecoding.jl package with a search radius of 10, a lateral variance penalty of 10, and no drops allowed. The simulation workflow is available here https://github.com/CaiGroup/UntanglingBarcodes/tree/main/simulations/Code_overlap_robustness.

### 3.14 Performance metrics for simulations

To evaluate the decoding performance in simulations, we calculate the sensitivity and FDR as

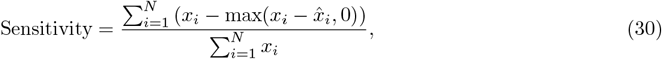

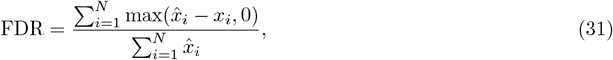

where *x*_*i*_ is the number of simulated transcript counts and 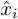 is the number of decoded transcript counts for the *i*th gene. *N* is the number of genes probed in the simulation.

### 3.15 Estimating sensitivity relative to smFISH in real data

For the sensitivity estimate of our method in cell culture, we compare the average number of barcodes of selected genes found per cell using our multiplexed seqFISH method to a comparison experiment that counts copies of mRNAs of the same genes using un-multiplexed sequential single-molecule FISH (smFISH) in a separate sample of cells from the same culture. The expression level distribution of the genes we chose to count with smFISH is representative of the expression level distribution of the entire panel probed using seqFISH (Supplementary Fig. 3B), We then fit a linear regression model of the average counts of mRNAs of each gene found per cell by seqFISH explained by each gene’s average mRNA count per cell as found by smFISH. The slope of the regression curve is the estimated sensitivity (Fig 2E).

### 3.16 Estimating overall false discovery rate in real data

To estimate the FDR in experiments, we decode each experiment twice: once using a codebook containing only codewords for which we designed probes to readout real transcripts and once using a codebook containing all of the codewords in the first codebook and all additional unused codewords of the same weight in the error-correcting code that are not probed for. These additional codewords serve as computational negative controls. Using two decoding runs improves the quality of the decoding results and the FDR estimate by eliminating competition from negative control codewords when decoding gene encoding codewords. We assume that negative control codewords are erroneously decoded at the same rate as gene encoding codewords on average (Supplementary Note 2). We estimate the average rate at which codewords are decoded erroneously as

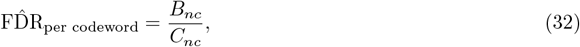

where *B*_*nc*_ is the number of negative control barcodes found when decoding including negative control codewords, and *C*_*nc*_ is the number of negative control codewords. We can estimate the number of falsely discovered gene encoding barcodes by multiplying the estimated false discovery rate per codeword, *B*_*nc*_*/C*_*nc*_, by the number of gene encoding codewords, *C*_*g*_. To estimate the proportion of gene encoding barcodes that are found in error (FDR), we divide the expected number of falsely discovered gene encoding barcodes by the total number of gene encoding barcodes found when decoding only gene encoding codewords, *B*_*g*_:

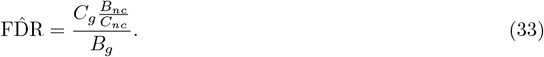

This equation has been used in previous works [15, 16, 20, 45].

### 3.17 Estimating FDR at different densities in real data

We estimate false discovery rates in real data at different local barcode densities using the same approach as for estimating the overall FDR, but partitioning the barcodes by local density. To find the local density of each barcode in each cell, we first construct a KDTree containing the spatial coordinates of barcodes found in the gene-encoding-only decoding run and a KDTree containing the negative control barcodes found for that cell. We then use each cell’s KDTree to find the number of barcodes within a 100 nm radius of each gene-encoding barcode found in the gene-encoding-only decoding run (including itself) and each negative control barcode found in the decoding run that included negative control barcodes. We then estimated FDR at each density using equation 33 with the number of gene encoding and negative control barcodes found at that density (Fig. 2D).

### 3.18 Primary-probe design

Gene-specific primary probes were designed as previously described with some modifications[9]. The probe length was set to be 33 nucleotides (nt), and a local BLAST query was run on each probe against the mouse transcriptome and a local database constructed from the probe sequences to ensure specificity. We managed to find 24 probes for the 271 selected highly expressed genes. We designed four secondary readout probe hybridization sites on each primary probe, with two readout probe hybridization sites on each end of the primary probe and the RNA-binding site in the middle. We selected 49 genes from the 271 genes for the smFISH experiment, with 36 probes per gene and two readout sites on each primary probe. The probe sequences are available as Supplementary Data.

### 3.19 Primary-probe construction

Primary probes were generated from oligoarray pools (Twist Bioscience) as previously described [9]. In brief, probe sequences were amplified from the oligonucleotide pools with limited two-step PCR cycles, and PCR products were purified using QIAquick PCR Purification Kit (Qiagen 28104). Then, in vitro transcription (NEB E2040S) was performed, followed by reverse transcription (Thermo Fisher EP0751). After reverse transcription, the single-stranded DNA (ssDNA) probes were alkaline hydrolyzed with 1 M NaOH at 65°C for 20 min to degrade the RNA templates and neutralized with 1 M acetic acid. Finally, probes were purified with SPRIselect beads (Beckman Coulter B23318) and eluted in nuclease-free water. Probe concentration was measured by Nanodrop.

### 3.20 Readout-probe design and synthesis

Readout probes of 15 nt in length were designed as previously described [9]. In brief, a set of probe sequences was randomly generated with combinations of A, T, G or C nucleotides. Readout-probe sequences within a GC-content range of 40–60% were selected. We performed a BLAST search against the mouse transcriptome to ensure the specificity of the readout probes. To minimize cross-hybridization of the readout probes, any probes with ten contiguously matching sequences between readout probes were removed. The reverse complements of these readout-probe sequences were included in the primary probes according to the designed barcodes. Readout probes of 15 nt in length were ordered from Integrated DNA Technologies as 5’-amine modified. The construction of readout probes was similar to that previously described [9]. In brief, 5 nmol DNA probes were mixed with 50 nmol of Alexa Fluor 647–NHS ester (A20006, Thermo Fisher) or Cy3B (PA63101, Cytiva) or Alexa Fluor 488–NHS ester (A30005, Thermo Fisher) in 0.5 M sodium bicarbonate buffer containing 5% dimethyl sulfoxide. The reaction was incubated at 37°C. After 1 hr, fresh 50 nmol of Alexa Fluor 647–NHS ester or Cy3B or Alexa Fluor 488–NHS ester was added to the solution. After another hour, 50 nmol of Alexa Fluor 647–NHS ester or Cy3B or Alexa Fluor 488–NHS ester was added a third time, making the final concentration of 3 mM in 0.5 M sodium bicarbonate buffer containing 15% dimethyl sulfoxide. Then, the DNA probes were subjected to ethanol precipitation, high-pressure liquid chromatography (HPLC) purification and column purification to remove all contaminants. Once resuspended in water, the readout probe concentration was quantified using Nanodrop and a 500 nM working stock was made. All the readout probes were stored at -20°C.

### 3.21 Coverslip functionalization

The coverslips were rinsed with 100% ethanol three times, and heat-dried in an oven at *>*90°C for 30 min. Then, the coverslips were cleaned with a plasma cleaner on a high setting (PDC-001, Harrick Plasma) for 5 min. Next, the coverslips were treated with 100 *µ*g *µ* l^−1^ of poly-d-lysine (P6407; Sigma) in water for *>*3 hrs at room temperature, followed by three rinses with water. The coverslips were then air-dried and kept at 4°C for no longer than 7 days.

### 3.22 Cell culture experiment

NIH/3T3 cells (ATCC) were cultured as previously described on the functionalized coverslips to ∼ 80–90% confluence [9]. Then, the cells were washed with 1× PBS once and fixed with freshly made 4% formaldehyde (28906, Thermo Fisher) in 1× PBS (AM9624, Invitrogen) at room temperature for 10 min. The fixed cells were permeabilized with 70% ethanol for 2 hrs at room temperature or overnight at -20°C. The cell samples were dried, and custom-made flow cells were attached to the coverslip. The samples were rinsed with three flow cell volumes (approximately 120 *µ*l) of 1X PBS and post-fixed with freshly made 7.5 mM BS-PEG5 (A35396, Thermo Fisher) in 1x PBS for 15 min at room temperature twice. Samples were rinsed with 3 flow cell volumes of 1X PBS to remove excess BS-PEG5. Afterwards, amines on the sample were blocked with 100 mM N-(Propionyloxy)succinimide (93535, Sigma) in 1x PBS. Samples were treated with 100mM N-(Propionyloxy)succinimide (NHS) in 1X PBS and 10% Dimethyl sulfoxide for 1 hrs at room temperature for three times (3 hrs total). The NHS solution was removed, and the samples were rinsed with 5 flow cell volumes (approximately 200 *µ*l) of 25% wash buffer (25% Formamide (4610, Sigma), 2x SSC, 0.1% Triton-X100 (93443-100ML, Sigma)). Hybridization buffer containing 0.1 mg/ml yeast tRNA (AM7119, Thermo Fisher) and 2 *µ*M polyTTG 200 nt (IDT) was added and the samples were incubated at 37°C for 1 hrs. Finally, primary probes (5 nM per probe for 24 probes per gene) were hybridized to the 271 genes in the hybridization buffer. The hybridization was allowed to proceed for 36 hrs in a humid chamber at 37°C. After hybridization, the samples were washed with 3 flow cell volumes of 30% wash buffer (30% Formamide, 2x SSC, 0.1% Triton-X100), followed by an incubation at 37°C for 30 min. Afterwards, the samples were rinsed with 6 flow cell volumes (approximately 240 *µ*l) of 4X SSC. Next, Tetraspeck beads (T7280, Thermo Fisher) were sonicated for 5 mins, diluted 1:100 in 4X SSC and vortexed. Tetraspeck beads were applied to the cell samples and incubated at room temperature for 5 min. Then, the sample was washed with 3 flow cell volumes of 1X PBS and post-fixed with 4% PFA in 1x PBS for 10 min at room temperature. Finally, the sample was washed with 6 flow-cell volumes of 4X SSC and sealed. The sample can be stored at 4°C for 2 weeks before imaging.

### 3.23 Mice

All animal care and experimental procedures were approved by the Caltech Office of Laboratory Animal Resources (OLAR). Wild-type C57BL/6J mice (8–12 weeks old; The Jackson Laboratory) were used for RS-code tissue experiments.

Mice were euthanized by CO_2_ inhalation, and brains were rapidly dissected following confirmation of death. Tissues were embedded in optimal cutting temperature (OCT) compound and snap-frozen in 2-methylbutane pre-cooled on dry ice. Embedded samples were stored at -80°C until cryosectioning.

### 3.24 Tissue slices experiment

Fresh-frozen mouse brain sections (10*µ*m) were fixed in 4% paraformaldehyde (PFA) in 1× PBS for 10 min at room temperature, washed with 1× PBS, dehydrated in 10% isopropanol for 1 min, air-dried and stored at -80°C. Before processing, sections were permeabilized in 70% ethanol overnight at 4°C, treated with 8% Triton X-100 in 1× PBS for 1h at room temperature, rinsed with 70% ethanol and air-dried. Flow cells were attached prior to subsequent processing.

Sections were post-fixed twice with 7.5 mM BS-PEG5 in 1× PBS for 15 min at room temperature. Free amines were blocked with 100 mM N-(Propionyloxy)succinimide in 1× PBS (4 × 45 min at room temperature), followed by five washes with 25% wash buffer (25% formamide, 2× SSC, 0.1% Triton X-100). Subsequent procedures were identical to those described for cell culture experiments, except that primary probe hybridization was performed for 48h at 37°C in a humidified chamber.

### 3.25 SeqFISH imaging for 3T3 cell culture

The imaging platform and automated fluidics delivery system were similar to those previously described with some modifications [9]. In brief, the flow cell on the sample was first connected to the automated fluidics system. Then the region of interest (ROI) was registered using nuclei signals stained with 30 *µ*g ml^−1^ DAPI (D8417; Sigma). Each serial hybridization buffer contained three unique sequences with different concentrations of 15-nt readouts conjugated to either Alexa Fluor 647 (50 nM), Cy3B (50 nM), or Alexa Fluor 488 (100 nM) in 10% hybridization buffer (10% formamide (4610; Sigma), 10% dextran sulfate (D4911; Sigma), and 4× SSC). The 150 *µ*l of serial hybridization buffers for 34 cycles of seqFISH imaging with a repeat for cycle 1 (in total 35 cycles) was pipetted into a 96-well plate. During each serial hybridization, the automated sampler moves to the well of the designated hybridization buffer and moves the 150 *µ*l hybridization buffer through a multichannel fluidic valve (EZ1213-820-4; IDEX Health & Science) to the flow cell (requires ∼ 40*µ*l) using a syringe pump (63133-01, Hamilton Company). The serial hybridization solution was incubated for 20 min at room temperature. After serial hybridization, the sample was washed with 500 *µ*l of 15% formamide wash buffer (15% formamide and 0.1% Triton X-100 in 2*×* SSC) to remove excess readout probes and non-specific binding. Then, the sample was rinsed with 500 *µ*l of 4*×* SSC, before being stained with DAPI solution (30 *µ*g ml^−1^ of DAPI in 4*×* SSC) for 2 mins. Next, an anti-bleaching buffer solution made of 10% (w/v) glucose, 1:100 diluted catalase (C3155, Sigma), 1 mg ml^−1^ glucose oxidase (G2133, Sigma), 5 mM Trolox (648471, Sigma), and 50 mM pH 8 Tris-HCl in 4× SSC was flowed through the samples. Imaging was done with a microscope (Leica DMi8) equipped with a confocal unit and lasers (Andor Dragonfly 200), a sCMOS camera (Andor Zyla 4.2 Plus), a 63× oil objective lens (Leica 1.40 NA) and a motorized stage (TMC 63-563). Snapshots were acquired with 0.25-*µ*m z-steps for 17 *z*-slices per field of view across 647-nm, 561-nm, 488-nm, and 405-nm fluorescent channels. Alexa Fluor 647, Cy3B, and Alexa Fluor 488 channels were acquired with 10% laser power and a 3,000-ms exposure time, while DAPI images were acquired with 10% laser power and a 200-ms exposure time. After imaging, stripping buffer (60% formamide and 0.1% Triton-X 100 in 2 *×* SSC) was flowed through for 1 min, followed by an incubation time of 3 min before rinsing with 4 *×* SSC solution. The serial hybridization, imaging and signal extinguishing steps were repeated for 34 cycles. Then, staining buffer for segmentation (50 nM LNA T20-Alexa Fluor 647 in 4 *×* SSC) was flowed in and allowed to incubate for 20 min at room temperature before imaging. Stripping buffer was applied to the sample to strip off the segmentation marker. Final blank images containing beads only were imaged as the last hybridization cycle. The integration of the automated fluidics delivery system and imaging was controlled by a custom-written script in *µ*Manager.

### 3.26 SeqFISH imaging for mouse brain slices

For mouse brain seqFISH imaging, images were acquired across 11 *z*-slices with 1-*µ*m z step. All 100 readout probes were conjugated to Cy3B and sequentially imaged over 100 hybridization cycles. Cy3B images were acquired with 10% laser power and a 3,000-ms exposure time. All other imaging and fluidics procedures were performed as described above for 3T3 cell culture seqFISH imaging.

### 3.27 smFISH experiment for 3T3 cell culture

smFISH experiments were performed as previously described [9]. The fixed cells were blocked and hybridized with the primary probes (10 nM per probe) in 25% hybridization buffer (25% formamide, 10% dextran sulfate and 4 × SSC) at 37°C for 24 hrs. The sample was washed with 30% wash buffer for 30 min at 37°C before imaging. The imaging platform was the same as the one in the seqFISH experiment. The serial hybridization, imaging, and signal extinguishing steps were repeated for 17 cycles. The sum of the gene counts per cell was analyzed using a previously published Python pipeline [20, 45].

