## Supplementary Information for "Untangling Overlapping Barcodes in Image-Based Spatial Genomics"

### 1 Supplement

#### 1.1 An explanation of the error-correcting codes in FISH-based spatial genomics

Linear error-correcting codes use codewords that are composed of sequences of symbols. Codewords can be divided into information symbols, which alone represent a message containing all the information encoded in a codeword, and parity check symbols, which are linear combinations of the information symbols that add redundancy and robustness to the codeword. The size of the alphabet from which the symbols are drawn is denoted as $q$, the number of symbols in a codeword is denoted as $n$, the number of information symbols is denoted as $k$, and the number of parity check symbols is denoted as $n-k$. The number of non-zero symbols in a codeword is called the weight of the codeword and is denoted as $w$.

In seqFISH, we conceptualize symbols as being transmitted from barcoded objects through a multicolored image where signals appear as dots and encode a symbol value determined by their color. In the first seqFISH experiments, each symbol was represented by a distinct fluorophore emitting a different color in the same hybridization [1, 2]. Later implementations of seqFISH expanded symbol block images to include pseudo-colors from images acquired in different hybridization cycles [3, 4]. This enabled experiments with expanded multiplexing capacity and reduced optical density by allowing larger alphabets ($q > 5$) from which symbols of the error-correcting code are drawn. SeqFISH, until now, has used $q$-ary parity check codes with $n = 4$ and $k = 3$ or $n = 5$ with $k = 3$ or 4 where all symbols in the alphabet, including zero, are represented by a pseudo-color. MERFISH uses weight-four codewords from binary error-correcting codes ($q = 2$) where zeros are represented by the absence of dots and are not probed. MERFISH has used Hamming codes [5] and non-linear codes [6].

One measure of an error-correcting code's robustness is its minimum Hamming distance between any pair of its codewords (MHD). Reed-Solomon codes are members of a special class of linear error-correcting codes, called maximum distance separable codes, for which MHD $= n - k + 1$. There are ways to increase the robustness of a code in ways that are not measured by the MHD. For example, Reed-Solomon codes use symbols from alphabets with size equal to a non-negative integer power of prime numbers. This enables using symbol blocks with multiple pseudo-colors, adding an extra rule that reduces ambiguity in superimposed barcodes: each barcode can only have one value-representing dot in each symbol block. Increasing the average Hamming distance between codewords without increasing the MHD of a code can also improve robustness. We find that thinning a codebook to not include all available codewords or adding parity checks that increase the average Hamming distance but not MHD can also increase the robustness of codes (Supplementary Fig. 2A). For linear error-correcting codes, the MHD is equal to the minimum weight of its non-zero codewords [7]. However, non-linear error-correcting codes, which do not have parity check symbols that are linear combinations of information symbols, are not subject to this constraint. This may provide an opportunity to encode information more efficiently and robustly at the cost of more computationally expensive decoding. To test the potential of non-linear error-correcting codes, we designed a non-linear error-correcting code. We first chose a minimum Hamming distance of 6, iterated over all combinations of 100 choose 5, and then added any combination to our codebook that is at least 6 Hamming distance away from any combination that had previously been added (Supplementary Fig. 2A). We expect that it is possible to construct better non-linear codes using more sophisticated discrete mathematics.

#### 2 Note on false discovery rate estimates

Our false discovery rate estimator, main text equation 33, relies on the assumption that, on average, negative control barcodes are decoded at the same rate per negative control codeword as gene encoding barcodes are erroneously decoded per gene encoding codeword. We have found that barcodes for some negative control codewords are found more often than others. However, we can explain most of the variation ($R^2 = 0.51$) in the decoding frequency of different negative control codewords using a linear model with explanatory variables constructed by counting the number of instances in which spatially overlapping gene encoding barcodes found in the decoding runs including only gene encoding barcodes recombine or nearly recombine into negative control barcodes and the total expression counts found of gene encoding barcodes in images where each negative control barcode is searched for (Supplementary Fig. 3C).

We now describe in detail how the model is constructed. $G$ and $C$ are binary representations of the codebook matrices for gene encoding and negative control codewords, respectively. $G$ is of dimension $n_g \times (n * (q-1))$ and $C$ is of dimension $n_c \times (n * (q-1))$, where $n_g$ is the number of gene encoding codewords, $n_c$ is the number of negative control codewords, $n$ is the number symbols in a codeword of the error-correcting code, $q$ is the size of the symbol alphabet for the error-correcting code, and $q-1$ is the number of pseudo-colors. Each row of these matrices represents a codeword. Each column corresponds to a readout image in the seqFISH experiment. Elements of $G$ and $C$, $s_{ij}$, are valued as

$$s_{ij} = \begin{cases} 1, & \text{if the seqFISH representation of the } i\text{th codeword includes a signal in the } j\text{th image} \\ 0, & \text{otherwise.} \end{cases} \tag{1}$$

We next define upper triangular overlap recombination matrices of dimension $n_g \times n_g$, $O(k)$, for each negative control codeword of arbitrary index $k$. Upper triangular elements of $O(k)$, $o_{ij}(k)$, count the number of dots needed to represent the $k$th negative control codeword that are available to recombine from spatial overlaps of the $i$th and $j$th gene encoding codewords

$$o_{ij}(k) = \begin{cases} (\mathbf{g_i} \vee \mathbf{g_j})\mathbf{c_k}', & \text{if } j > i \\ 0, & \text{otherwise,} \end{cases} \tag{2}$$

where $\mathbf{g_i}$ and $\mathbf{g_j}$ are the $i$th and $j$th row vectors of $G$, the binary representations of the $i$th and $j$th gene encoding codewords, respectively. $\mathbf{c_k}$ is the $k$th row vector of $C$, the $k$th binary negative control codeword vector, and $\vee$ is the element-wise logical or operator.

We construct recombination frequency matrices of dimension $n_g \times n_g$, $P(r)$, that hold the number of co-occurrences of gene encoding barcodes of different kinds within a search radius, $r$. Elements of $P(r)$, $p_{ij}(r)$, are equal to the number of instances in the entire dataset in which barcodes of the $i$th codeword are located within a spatial distance $r$ of the barcodes of the $j$th codeword. To aid us in writing this definition formally, let be $\mathbf{u_{ik}}$ the position vector of the $k$th barcode of the $i$th gene $\mathbf{v_{jl}}$ is the position vector of the $l$th barcode of the $j$th negative control codeword. Then,

$$p_{ij}(r) = \begin{cases} \text{sum}\left(\left\{\delta(||\mathbf{u_{ik}} - \mathbf{v_{jl}}||_2 \leq r)\middle| k \text{ in } \{1,2,...,n_i\}, l \text{ in } \{1,2,...,n_j\}\right\}\right), & \text{if } j > i \\ 0, & \text{otherwise,} \end{cases} \tag{3}$$

where $n_i$ and $n_j$ are the number of barcodes that were decoded for the $i$th and $j$th codewords respectively. Now we can define the vector-valued function

$$\boldsymbol{\rho}(n_\rho, r) = [\rho_1(n_\rho, r), \rho_2(n_\rho, r), ..., \rho_k(n_\rho, r), ..., \rho_{n_c}(n_\rho r)], \quad (4)$$

that generates explanatory variables of length $n_c$ for our linear model. The $k$th element of $\boldsymbol{\rho}(n_\rho, r)$ is calculated as

$$\rho_k(n_\rho, r) = \text{sum}\Big(P(r) \odot \big(O(k) == n_\rho\big)\Big), \quad (5)$$

where $n_\rho$ is the desired number of dots from a target codeword found in an overlap, $r$ is the search radius within which barcodes are considered overlapping, == is the element-wise logical equal to operator, and $\odot$ is the element-wise multiplication operator (Hadamard product operator). Our linear model predicts the frequencies at which barcodes for each negative control codeword, $\mathbf{d_c}$, are found as

$$\hat{\mathbf{d}}_\mathbf{c} = \beta_1 + \beta_2(CG'\mathbf{d_g}) + \beta_3\boldsymbol{\rho}(3,1) + \beta_4\boldsymbol{\rho}(4,1) + \beta_5\boldsymbol{\rho}(3,2) + \beta_6\boldsymbol{\rho}(4,2), \quad (6)$$

where $CG'\mathbf{d_g}$ gives the exposure of each negative control codeword to dots transmitted by gene encoding codewords encoded in the same images, $\mathbf{d_g}$ is the vector containing the number barcodes for each gene that were found in the decoding run that considered only gene encoding barcodes, $\beta_1$ is the intercept and $\beta_2$ to $\beta_6$ are fit coefficients.

Our model does not include possible effects from registration errors (Supplementary Fig. 5D), which are unevenly distributed in readout images because they are different in each field of view and would require a much more complicated model to account for. Other factors that may contribute to variation in the frequency at which different negative control barcodes are decoded may include non-specific binding and failure to distinguish spatially overlapping dots read out from two different molecules in the same image. It is likely that negative control barcodes are found in error more often than gene encoding barcodes because we construct codebooks to maximize the distance between gene encoding barcodes, meaning that gene encoding barcodes are on average more similar to negative control barcodes than to each other.

#### 2.1 SeqFISH+ simulations and limits on resolving dense barcodes from image fitting

##### 2.1.1 Spots-first simulations

The fitting algorithm we use in our spots-first decoding workflow, ADCG, cannot effectively resolve separate point spread functions in the same optical image centered within a full width at half maximum of each other. Partially overlapping dots in the same image may also cause fitting errors, resulting in fit dots whose coordinates do not correspond to those of the actual molecules. To investigate the impact of errors in image fitting on decoding, we simulated synthetic image data and dot location data modeled to approximate the conditions of the previously published 10,000 gene seqFISH+ experiment [3]. In these simulations, not only were simulated dot locations fed directly into the decoder, but synthetic images were produced as a linear combination of Gaussian point spread functions centered at each simulated dot location. Images were fit with ADCG[8], then the fit dot locations recovered from the image fitting were decoded. We simulated 70 replicates of seqFISH+ data in a 5×5 pixel ROI with various densities of barcodes and non-specific dots with locations drawn from the uniform distribution. The $x$ and $y$-coordinates of dots read out from each barcode are drawn from a normal distribution with a standard deviation of 0.5 pixels around the simulated true location of the barcode. Simulated images are generated as linear combinations of Gaussian point spread functions with a sigma of 1.2 pixels and the same brightness from both barcodes and non-specific binding. ADCG attempts to recover the dot locations from the simulated images, and both the ADCG recovered dots and the directly simulated dot locations are decoded by syndrome decoding with a position variance penalty of 6 and a log brightness variance penalty of 0. We found a decrease in decoding sensitivity and increased robustness to extra non-specific dots in the image-based simulations compared to the image-less simulations (Supplementary Fig. 6A). Performance metrics for the simulation were calculated as described in the methods section for simulations.

##### 2.1.2 Resolving overlapping barcodes in single images using regulated regression

The spots-first approach to resolving ambiguity arising from overlapping barcodes only addresses cases in which the overlapping barcodes are not probed in any of the same images. To further increase the density at which we can resolve barcodes, we will need to be able to handle situations when two spatially overlapping barcodes are probed in the same image. It has previously been shown that a compressed sensing approach can resolve overlapping dots in STORM images [9]. Others have used a similar approach to fit sparse models of spatial genomics image data as generated by barcode-specific source functions using $l_1$-regularization [10]. Here, we show in simulation that this idea can be extended to resolving many imperfectly aligned overlapping barcodes by first identifying candidate barcodes to use as explanatory variables and then using more stringent $l_0$-regulated regression.

We simulated image data where the location $(x_i, y_i)$ of the $i$th barcode is drawn from a 2D uniform distribution between 0 and 5. Synthetic images are 9×9 pixels. Readout point spread functions with sigma of one from the $j$th dot of each $i$th barcode are added to the synthetic images in which they are expected and centered at $x_{ij} = x_i + 2 + \epsilon_{x_{ij}}$ and $y_{ij} = y_i + 2 + \epsilon_{xij}$. The addition of 2 to $x_i$ and $y_i$ serves to pad the edges of the ROI, ameliorating edge effects. The localization errors are drawn as $\epsilon_{xij} \sim N(x_{ij}, 0.2)$ and $\epsilon_{yij} \sim N(y_{ij}, 0.2)$. The first step in the decoding process is to find candidate locations using the same method described in the compressed sensing decoding subsection of the methods section. Next, candidate barcodes are identified by syndrome decoding with a search radius of 0.2 pixels.

We next use an $l_0$-regulated regression model to choose the final set of barcodes from the candidate list,

$$\hat{\boldsymbol{\beta}} = \text{argmin} \frac{1}{2} ||\mathbf{y} - A_b \boldsymbol{\beta} - \beta_0||_2^2 + \lambda ||\boldsymbol{\beta}||_0. \tag{7}$$

This model is nearly identical to equation 27 of the main text. The only differences are that it uses $l_0$ regularization instead of $l_1$ regularization, there are no differential penalties for different non-zero elements of $\boldsymbol{\beta}$, and the normalization factor of the sum squared errors term (decided by the L0learn package)[11, 12]. Using this procedure, we were able to nearly perfectly decode dense simulated images encoded using the same Reed-Solomon code experimentally implemented in this manuscript (Supplementary Fig. 6B) without a duplicate removal step that is needed following optimizing with equation 27 of the main text.

#### 3 Supplemental Figures

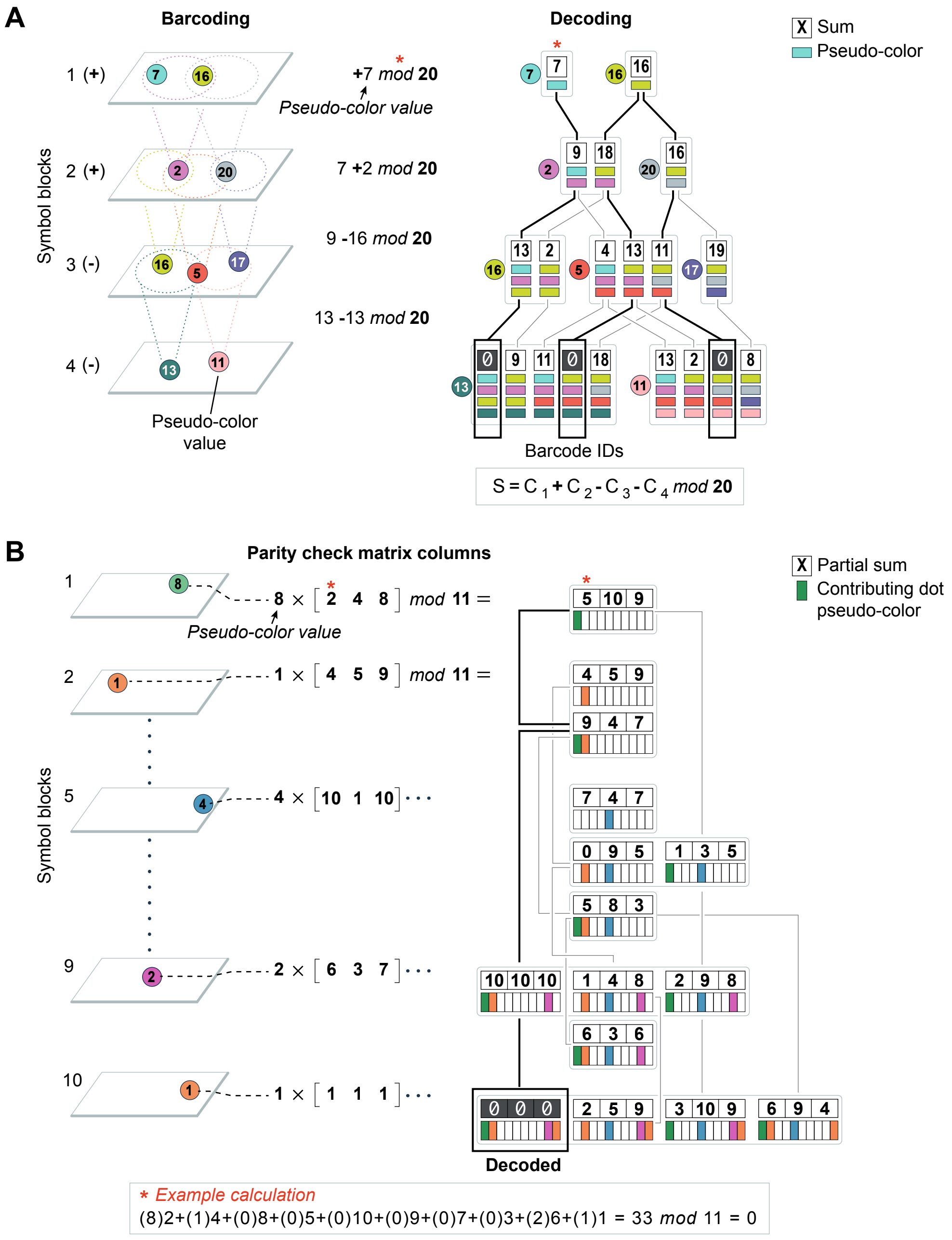


**Supplementary Figure 1**

**Supplementary Figure 1** A) We illustrate the simplified dynamic programming algorithm for evaluating syndromes in the $q$-ary parity check seqFISH code used in the published 10,000 gene seqFISH+ experiment. Candidate barcodes are identified by summing syndromes from numbered pseudo-colors of dots that align in subsequent symbol blocks. Dots in the first symbol block are assigned arrays holding their pseudo-color numbers. Each dot in the second symbol block is assigned an array found by adding its pseudo-color number element-wise to the concatenation of partial sum arrays of dots in the first symbol block that align within a search radius. This procedure is repeated for dots in the third and fourth symbol blocks, but subtracting their pseudo-color number. Sums evaluating to zero in the fourth symbol block indicate valid barcodes where contributing dots are traced. B) We illustrate a more general case of our dynamic programming algorithm for syndrome decoding with the error-correcting code used in our new experiment. This case differs from the case in panel A in two ways. First, three parity check equations are summed, so three partial sums are stored where a single partial sum would be stored in panel A. Second, zero-valued symbols are not probed and are represented by the absence of a dot in a symbol block. Consequently, the dots are aligned to neighbors in multiple preceding symbol blocks and the number of dots contributing to each partial sum is tracked. The dots in the second through $n - w + 1$st symbol block must search for aligning neighbors in all previous symbol blocks. Dots in symbol blocks $b \in [n - (w - 2), n]$ must only continue adding to partial sums with neighboring dots containing partial sums of $n - b$ or more dots since $w$ dots are needed to form a barcode.

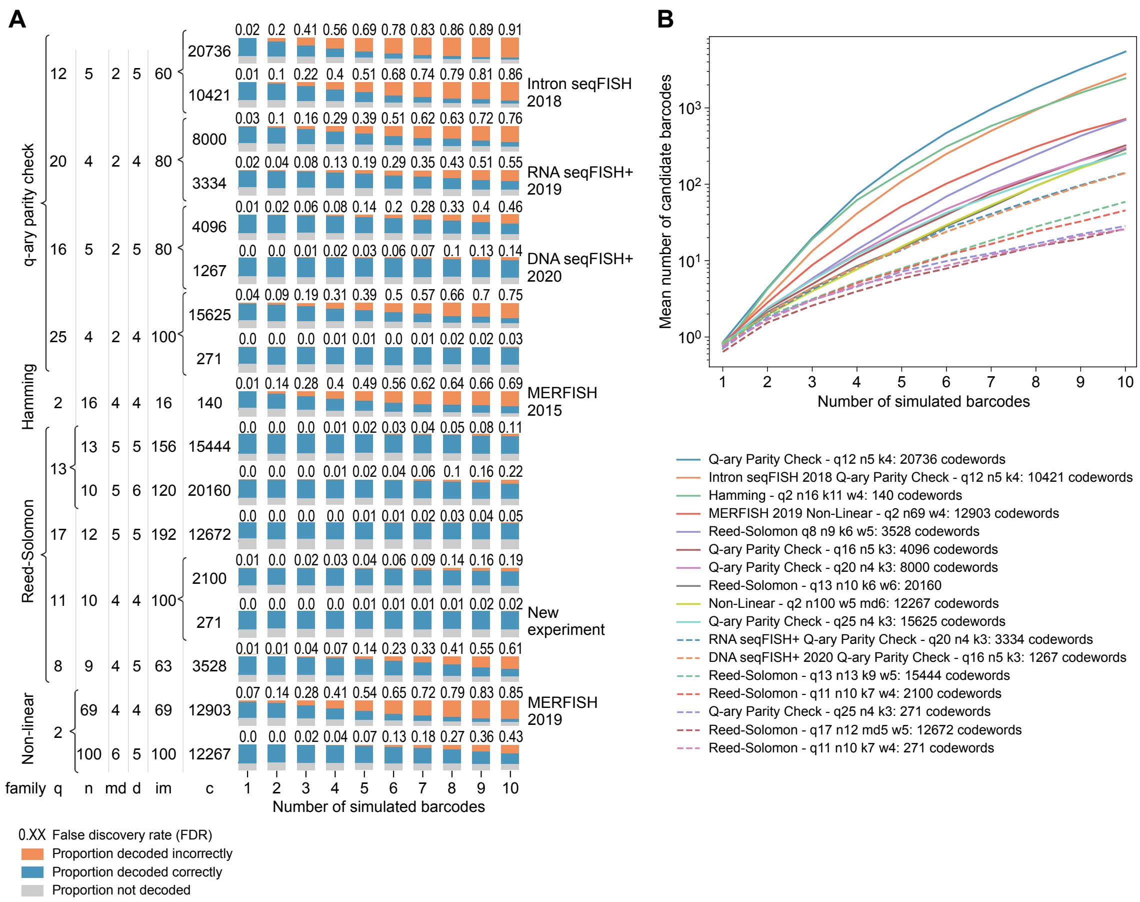


**Supplementary Figure 2** A) Expanded version of main Fig. 1E. The first eight rows show simulations for codes from the $q$-ary parity check code family used in previous seqFISH experiments. Brackets denote branching families of related codes. The first $q$-ary parity check code in each bracket pair uses all codewords from the code of the given weight. The second uses a subset of the codewords in the first that were used experimentally in this or a previous paper, except for the 271 codeword subset of the q25n4k3 single parity check code, which matches the experimental characteristics of the Reed-Solomon encoded experiment in this work. B) A plot of the mean number of candidate barcodes arising in simulations of various codes as a function of the number of true barcodes in the simulations. These results are drawn from the same simulations shown in panel A. Codes that produce fewer spurious candidate solutions in the simulations have higher decoding sensitivity and lower false discovery rates in the same simulations than codes that produce more spurious candidate solutions.

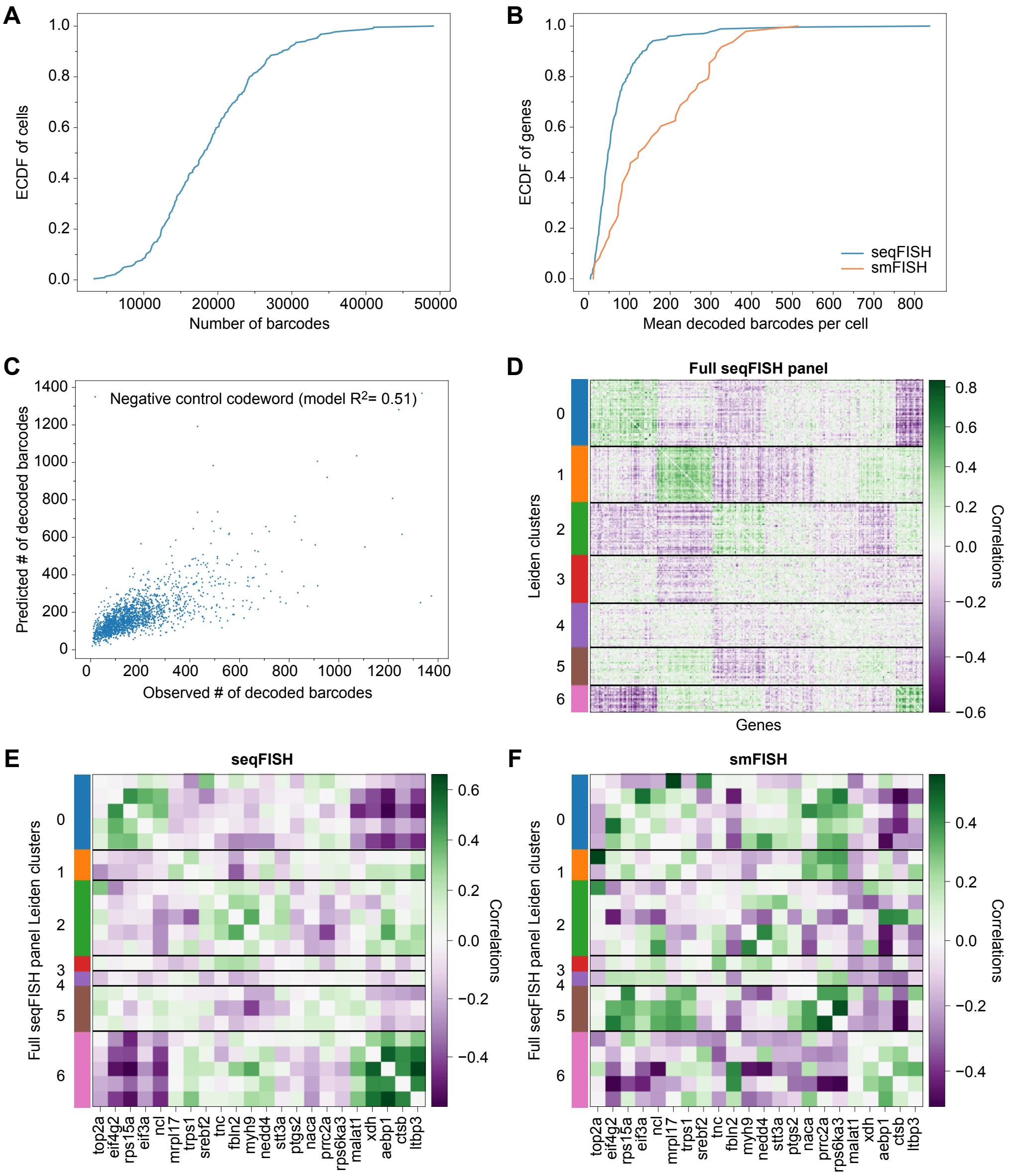


**Supplementary Figure 3** Summary plots for the Reed-Solomon encoded cross-channel NIH 3T3 cell experiment. A) An empirical cumulative distribution function of the number of gene encoding barcodes decoded in each cell of the cross-channel Reed-Solomon encoded seqFISH using decoding parameters for which summary statistics are reported in the text: lateral variance penalty $= 3$ and one drop allowed. B) The distribution of the mean barcodes per cell found for genes in the multiplexed seqFISH experiment compared with the un-multiplexed sequential smFISH experiment. The distribution for the seqFISH counts is approximately the same as the distribution for the sequential smFISH counts multiplied by the estimated sensitivity, 0.49 (Fig. 2E). C) Barcodes for different negative control barcodes are found at different frequencies. We can explain most of the variation using a linear model that uses variables constructed from the list of gene encoding barcodes per-cell encoded in the gene encoding only decoding run. D) Per-cell gene-gene correlations in our 271 gene 3T3 cell experiment. There are seven Leiden clusters of genes that are correlated with each other and anti-correlated with genes of at least one other cluster. E-F) Panel E shows a subset of the per-cell gene-gene correlation matrix shown in panel D, which is similar to the per-cell gene-gene correlation matrix found using the reference smFISH measurements of the same genes (F).

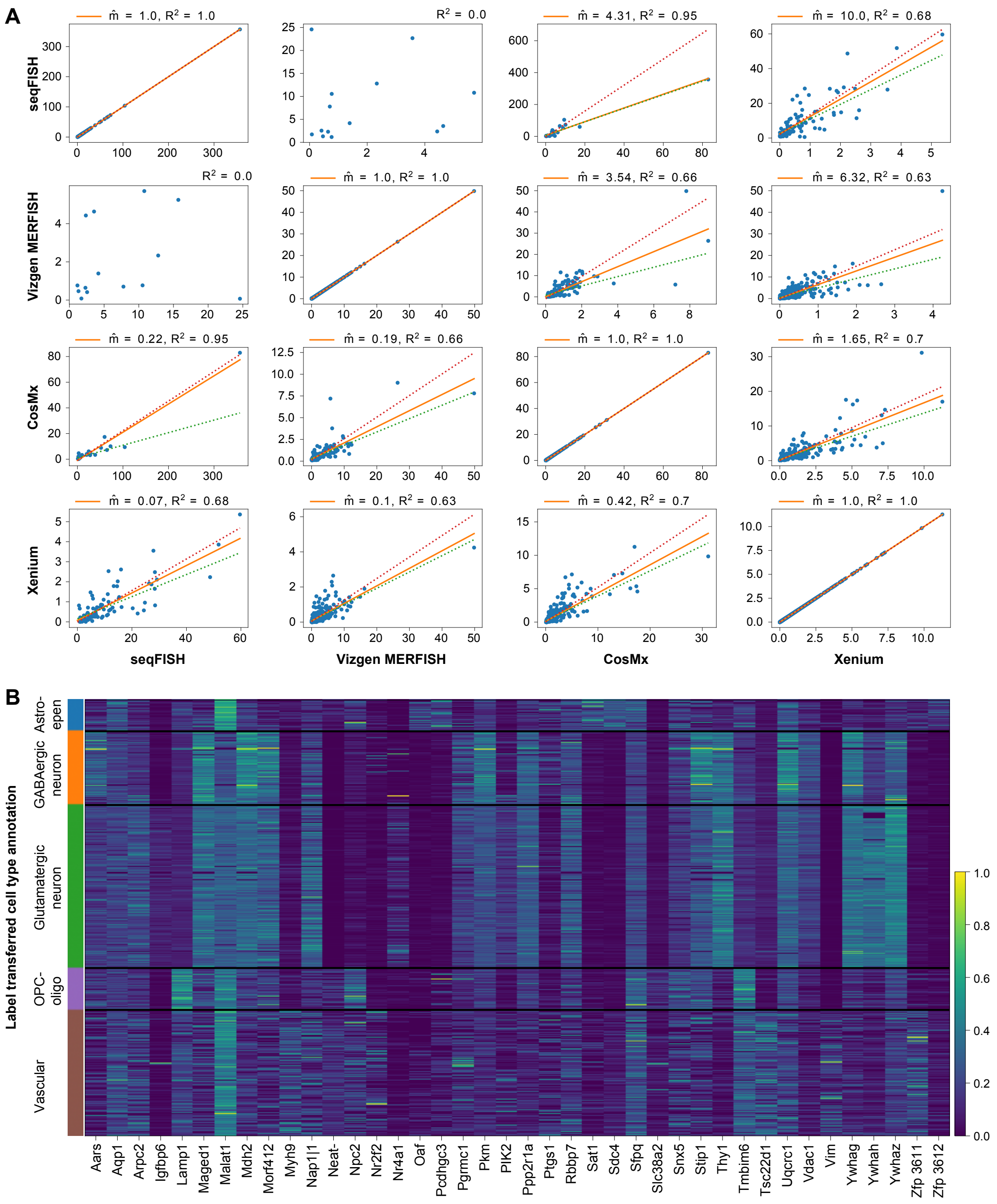


**Supplementary Figure 4** A) Correlations and relative sensitivities between our method and selected commercial imaging-based spatial genomics methods when applied to the mouse brain. $\hat{m}$ is the predicted slope (relative sensitivity) of the best-fit model using all genes measured by both methods. Dotted lines show the upper (red) and lower (green) bounds of the 95% confidence interval of the slope found using a non-parametric bootstrap. There were too few genes measured in both our seqFISH experiment and the Vizgen sample dataset to find a model with a correlation. B) Gene by cell matrix heatmap for mouse cortex cells profiled by seqFISH, including only cells whose cell type was predicted by a consensus of CONCORD models trained on different balanced subsets of Allen Brain Atlas 10X single cell sequencing data.

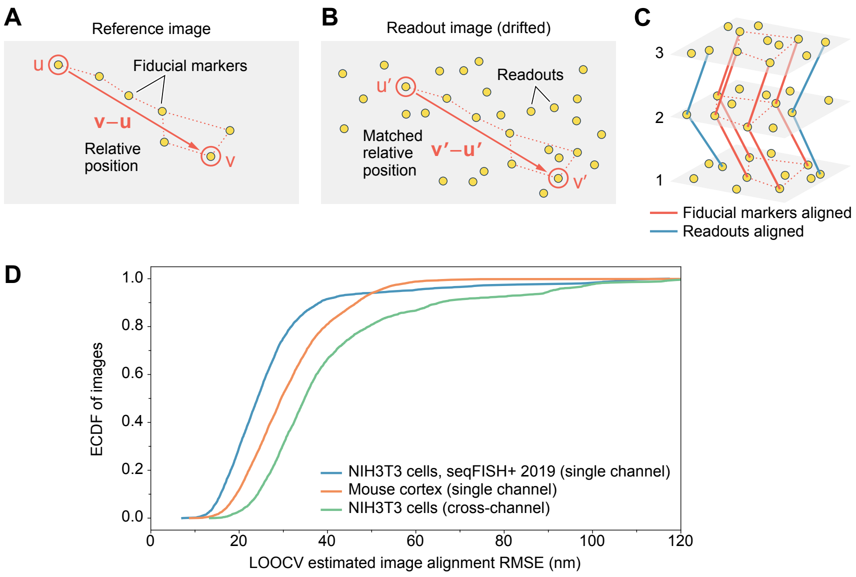


**Supplementary Figure 5** We register images using diffraction-limited objects as fiducial markers. A) Reference images of the fiducial markers alone show a constellation of fiducial markers. Each pair of fiducial markers, $u$ and $v$, have a known relative position from each other, $\mathbf{v} - \mathbf{u}$, all of which together describe the shape of the fiducial marker constellation mathematically. B) In readout images, the dots from the fiducial marker constellation are still visible, but there are also additional dots read out from target molecules. To find the constellation of fiducial marker dots in readout images, we first search for a pair of dots, $u'$ and $v'$, at approximately the same relative position to each other as a distant pair of fiducial markers, $u$ and $v$, are to each other in the reference image ($||(\mathbf{v}-\mathbf{u})-(\mathbf{v'}-\mathbf{u'})||_2 \leq \epsilon$). After finding a matching $u'$ and $v'$ at the same relative position as $u$ and $v$, we can find the remaining fiducial markers in the fiducial marker constellation at their expected relative locations to $u'$ and $v'$ in the readout image. C) We correct drifts and chromatic aberration in readout images by aligning the matched fiducial markers (red paths) to align signals from barcoded molecules (blue paths). D) We estimated the alignment error attained using fiducial marker matching registration with leave-one-out cross-validation in each of the three datasets we analyzed. Alignment accuracy had a median RMSE of 24 nm in single-channel encoded 3T3 cell culture, 29 nm in mouse cortex (for fields of view that were not discarded), and 35 nm for cross-channel encoded 3T3 cell culture.

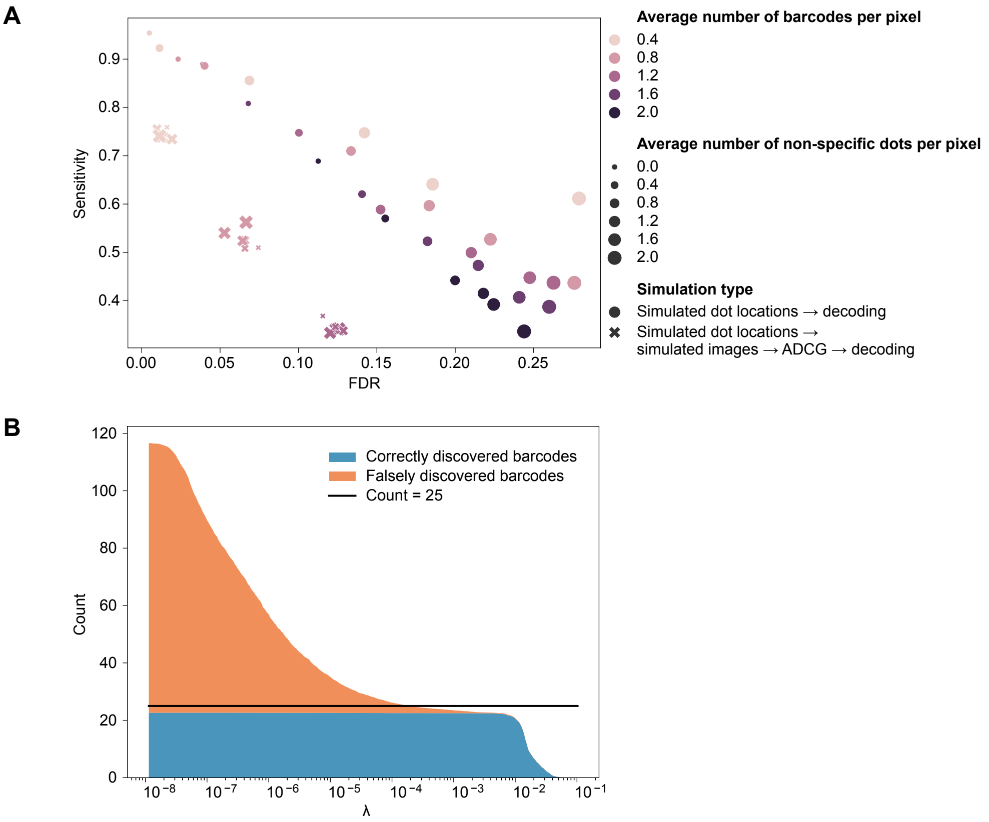


**Supplementary Figure 6** A) Decoding performance on simulated seqFISH+ images and dot locations with various barcode and non-specific dot densities. Barcode and non-specific dot densities are for the entire simulated experiment, not individual images. B) Decoding by regulated regression can distinguish overlapping barcodes in simulated images. This plot shows the average performance of 100 simulation replicates of 25 barcodes in a simulated 5x5 pixel (500x500 nm) area with 2 pixels of padding in the synthetic images to ameliorate edge effects. The x-axis varies the regularization parameter, $\lambda$, in the final L0-regularized regression to choose barcodes that explain the simulated data. In these simulations, the optimal value of the regularization parameter is around 0.005. Lower regularization parameters lead to overfitting and return excessive false solutions.
